# Extra-field activity shifts the place field center of mass to encode aversion experience

**DOI:** 10.1101/218180

**Authors:** Omar Mamad, Beshoy Agayby, Lars Stumpp, Richard B. Reilly, Marian Tsanov

## Abstract

The hippocampal place cells encode spatial representation but it remains unclear how they store concurrent positive or negative experiences. Here, we report on place field reconfiguration in response to an innately aversive odor trimethylthiazoline (TMT). The advantage of TMT is the absence of learning curve required for associative fear conditioning. Our study investigated if CA1 place cells, recorded from behaving rats, remap randomly or if their reconfiguration depends on the location of the aversive stimulus perception. Exposure to TMT increased the amplitude of hippocampal beta oscillations in two arms of a maze (TMT arms). We found that a population of place cells with fields located outside the TMT arms increased their activity (extra-field spiking) in the TMT arms during the aversive episodes. These cells exhibited significant shift of the center of mass towards the TMT arms in the subsequent post-TMT recording. The induction of extra-field plasticity was mediated by the basolateral amygdala complex (BLA). Photostimulation of the BLA triggered aversive behavior, synchronized response of hippocampal local field oscillations, augmented theta rhythm amplitude and increased the spiking of place cells for the first 100ms after the light delivery. This occurred only for the extra-field-but not for intra-field spikes. Optogenetic BLA-triggered an increase in extra-field spiking activity correlated to the degree of place field plasticity in the post-ChR2 recording session. Our findings demonstrate that the increased extra-field activity during aversive episodes mediates the degree of subsequent field plasticity.

## Introduction

Hippocampus temporally encodes representations of spatial context-dependent experiences (Knierim, 2003) and these memory traces are functionally strengthened in the cortical areas for long-term recollection (Nadel and Moscovitch, 1997; Kitamura et al., 2009; Tayler et al., 2013; Denny et al., 2014). Current theories propose that memory of spatial location is encoded by hippocampal place cells (O’Keefe and Nadel, 1978), but there is scarce information how these neurons encode non-spatial information such as aversive episodes. We know that aversion evokes place field remapping (Moita et al., 2004; Kim et al., 2015) where a subset of neurons in hippocampal area CA1 change their preferred firing locations in response to predator odor (Wang et al., 2012). However, it is still unclear which neurons remap to encode fearful experience and which neurons preserve their spatial fields. Here, we examined the principles governing aversion-induced place field remapping. We tested the hypothesis that the place cells remapping depends on the spatial location of the aversive stimulus perception. The evaluation of change of place field center of mass (ΔCOM) is the most sensitive indicator of experience-dependent place field place field reconfiguration (Mehta et al., 1997; Knierim, 2002; Lee et al., 2004; Lee et al., 2006). We evaluated here the aversion-induced long-term shift of the ΔCOM for all place fields. Beta frequency band (15-30Hz) has been reported as a reliable indicator for the detection of aversive olfactory signals by the limbic circuitry (Igarashi et al., 2014). We analyzed the amplitude of hippocampal beta frequency to determine which section of the maze was the main location of the aversion odor perception (Trimethylthiazoline - TMT).

The amygdala supports aversive associative memories and relates adaptive behavior to the emotional valence of sensory stimuli (Phelps and LeDoux, 2005; Sadrian and Wilson, 2015). Amygdalar signaling of aversion controls the stability of hippocampal place cells and lesioning of the amygdala prevents the effect of fearful experience on the place fields remapping (Kim et al., 2015). Inactivation of amygdala blocks tone-induced fear conditioning that triggers place field remapping (Donzis et al., 2013), while amygdalar activation reduces the field stability of hippocampal place cells (Kim et al., 2012). Aversion-induced activation of the pyramidal neurons in amygdala mediates the formation of spatial context-dependent place aversion and in parallel with new hippocampal engrams (Ramirez et al., 2013; Ryan et al., 2015). However, there have been no direct tests of how new hippocampal ensembles are formed. To understand the mechanism of aversion-induced population response from hippocampal spatial representation we optogenetically activated pyramidal neurons from basolateral complex of amygdala (BLA). We photostimulated BLA to determine if the patterns of ensembles reconfiguration after aversion-induced place field remapping are replicated after optogenetic BLA activation. We investigated if the degree of place field plasticity differs when the spatial optogenetic stimulation was applied to the main place field compared to optogenetic stimulation of the extra-field activity. Extra-field spikes, previously considered to be noise, are now proposed to play an essential role in information processing, learning and memory formation (Johnson and Redish, 2007; Johnson et al., 2009; Epsztein et al., 2011; Ferguson et al., 2011; Wu et al., 2017). Here, we explored if intra- or extra-field spikes mediate experience-dependent encoding of aversive episodes.

## Materials and Methods

### Ethics Statement

We conducted our experiments in accordance with directive 2010/63/EU of the European Parliament and of the council of 22 September 2010 on the protection of animals used for scientific purposes and the S.I. No. 543 of 2012, and followed Bioresources Ethics Committee (individual authorization number AE19136/I037; Procedure Numbers 230113-1001, 230113-1002, 230113-1003, 230113-1004 and 230113-1005) and international guidelines of good practice (project authorization number: AE19136/P003).

### Animals

Male, 3-6 months old, Lister-Hooded rats were individually housed for at least 7 days before all experiments, under a 12-h light–dark cycle, provided with water *ad libitum*. Prior the experiments restricted feeding diet kept the rats on 80% of their expected weight when fed ad libitum. Experiments were performed during the light phase.

### Surgical implantation of recording electrodes and recording techniques

Eight tetrodes and optic fiber were implanted in hippocampal CA1 area: −3.8 AP, 2.3 ML and 1.8 mm dorsoventral to dura. The optic fiber and tetrodes were implanted unilaterally in BLA: 2.4 AP, 4.9 ML, and 7.0 mm dorsoventral to dura. After a minimum 1 week recovery, subjects were connected, via a 32-channel headstage (Axona Ltd.) to a recording system, which allowed simultaneous animal position tracking. Signals were amplified (10000- to 30000-fold) and band-pass filtered between 380 Hz and 6 kHz for single-unit detection. To maximize cell separation, only waveforms of sufficient amplitude (at least three times the noise threshold) were recorded. Candidate waveforms were discriminated off-line using graphical software (Tint, Axona Ltd.), which allows waveform separation based on multiple features including spike amplitude, spike duration, maximum and minimum spike voltage, and the time of occurrence of maximum and minimum spike voltages. Autocorrelation histograms were calculated for each unit, and the unit was removed from further analysis if the histogram presented spiking within the first 1 ms (refractory period), inconsistent with good unit isolation. Only stable recordings across consecutive days were further analyzed. The stability of the signal was evaluated by the cross-correlation of spike amplitudes and similarity comparison of the spike clusters between the sessions and cluster distributions. The single unit signals from the last recording session and the probe were compared for waveform similarity, cluster location, size, and boundaries. Peak and trough amplitudes of the averaged spike waveforms were compared using Pearson’s *r*. Values for *r* ≥ 0.8 indicated that the same populations of cells were recorded throughout the last recording session and the probe.

### Hippocampal unit identification and spatial firing analysis

Single hippocampal pyramidal cells and interneurons were identified using spike shape and firing frequency characteristics (Ranck, 1973; Wilson and McNaughton, 1993). Firing rate maps allow for visual inspection of neurons preferred areas of firing (i.e. place fields). They were constructed by normalizing the number of spikes which occurred in specific pixelated coordinates by the total trial time the animal spent in that pixel. This produced maps depicting the place fields of each cell. Maps were quantified in Hz (smoothed maps). We defined place field size as the region of the arena in which the firing rate of the place cell was greater than 20% of the maximum firing frequency (Brun et al., 2002). Appearance of sharp waves and ripples during immobility, triggers the spiking of multiple place cells (Wu et al., 2017). To avoid spikes reactivation during sharp wave ripple we excluded spikes from epochs with running speeds below 5 cm/s (Alme et al., 2014; Grosmark and Buzsaki, 2016). The place field analysis included only epochs during which the animal’s velocity was at least 5 cm/s.

### Extra-field spiking

Place fields were defined as areas of 9 contiguous pixels (2.5 cm^2^ / pixel) with average activity >20% of the field maximum rate. Extra-field spiking was defined as spikes occurring outside of the identified place field areas (Huxter et al., 2003; Johnson and Redish, 2007). The extra-field spiking thus included secondary place fields with sizes smaller than 9 contiguous pixels or with averaged firing rate smaller than 20% of the maximum firing rate (Brun et al., 2002; Huxter et al., 2003; Johnson and Redish, 2007).

### Measurement of local field activity

The local field potential (LFP) was sampled at 250 Hz and stored for further off-line analysis. LFP signal frequency analysis was carried out using MATLAB’s Signal Processing Toolbox (MATLAB, Natick, MA) where the power was calculated using the Short-Time Fourier Transform of the signal (Hanning window of 2s, with overlap of 1s) and interpolated into color-coded power spectrograms. Information was displayed as the magnitude of the time-dependent Fourier Transform versus time in a color gradient graph with the maximum corresponding to 0 dB.

### Phase-locking value

To evaluate the effect of optogenetic BLA stimulation we compared the hippocampal local field oscillations of a single electrode between multiple trials (Mamad et al., 2015). Phase-locking statistics measures the significance of the phase covariance between separate signals and allows direct quantification of frequency-specific synchronization (i.e., transient phase-locking) between local field potentials (Lachaux et al., 1999). The phase-locking value is the amplitude of the first circular moment of the measured phase difference between two phases (Lachaux et al., 1999; Canolty et al., 2010). The phase-locking value ranges between 0 and 1; 0 signifying purely random rise and fall whereas a value of 1 signifies that one signal perfectly follows the other. To distinguish between noise-related fluctuations of the phase-locking values we compared the observed data with shuffled data (Mamad et al., 2015).

### Experimental design

The animals were trained to navigate between the northwest and southeast corners **of** rectangular-shaped linear track, where two pellets were continuously positioned. The animals were allowed to freely navigate in both clockwise and counter-clockwise directions of this rectangular-shaped linear track (10cm width, 85cm length of the arms): via the southwest arms and via the northeast arms. For the TMT experiments one of the filter papers of the track was scented with 50 µl 10% trimethylthiazoline (Contech). The advantage of trimethylthiazoline (TMT) is the absence of learning curve required for associative fear conditioning. The experimental protocol involving one TMT session with duration of 12 min was designed to evoke long-lasting (>24 hours) but weak place aversion response during the post-TMT recording session. The place aversion was measured only in the first 60 seconds of the post-TMT recording session. In the subsequent 11 minutes of the post-TMT recording session the animals displayed regular navigation in the TMT arms. This protocol allowed for sufficient number of passes, preventing an undersampling path measurement of the post-TMT navigation. During the TMT sessions the TMT filter papers were applied in all locations of the TMT arms for different rats; this protocol was designed to match the ChR2 protocol where the blue light was applied across all locations of the ChR2 arms. For the optic stimulation sessions, the laser was switched on when the animal entered the south arm or the west arm with continuous photostimulation trains (473 nm, 50 Hz, 5 ms pulse duration, 12 pulses per train, 0.5 sec inter-train interval) until the animal exited this section of the track. The blue laser was synchronized with the video-tracking and with the recording system through hardware and DACQBASIC scripts (Axona. Ltd). The duration of each session (baseline, TMT, ChR2) was 12 min. The TMT zone (ChR2 zone) included the TMT arms (ChR2 arms) and the feeding corners. For control experiments we used filter papers of the track scented with 50 µl 10% ethanol, which was a familiar odor to the rats. The animals were habituated prior the recording sessions to the scent of ethanol.

### Clockwise and counter-clockwise place field analyses

The rat’s direction of movement was calculated for each tracker sample from the projection of the relative position of the LEDs onto the horizontal plane. The momentary angular displacement was calculated as the difference in the animal’s position between successive 50 Hz time samples. The direction time series was first smoothed by calculating a five-point running average. After smoothing, the instantaneous direction of movement was calculated as the angular displacement between successive points per time (Taube, 1995). To restrict the influence of inhomogeneous sampling on directional tuning, we separated the directionality for the pre-TMT(ChR2) and post pre-TMT(ChR2) where the animals exhibited consistent navigation, but not for the TMT(ChR2) sessions where the direction of animal’s navigation was highly inconsistent due to the aversive episode. For the linear color-coded representation of the unidirectional place fields the firing rate was normalized for each cell to the cell’s baseline maximal firing rate. The unidirectional clockwise / counter-clockwise place fields were defined as areas of 9 contiguous pixels (2.5 cm^2^ / pixel) with average activity >10% of the field maximum rate. Although the reduction of the place field firing rate cut-off to 10% increases the extra-field noise this approach also preserves the peripheral spiking activity considered outside the place field with the 20% cut-off approach. The firing rate difference between the pre-TMT(ChR2) and post-TMT(ChR2) recordings was normalized by the ratio of the difference over the sum of the pre- and post-spiking count (see spike ratios).

### Optogenetic tools

AAV-CaMKIIa-hChR2(H134R)-eYFP-WPRE-hGH viral construct was serotyped with AAV5 coat proteins and packaged by Vector Core at the University of North Carolina with viral titers ranged from 1.5-8 x 10^12^ particles per mL. For control experiments we used virus bearing only the YFP reporter. Randomization of group allocation (ChR2 versus YFP controls) was performed using an online randomization algorithm (http://www.randomization.com/). The virus injection was applied unilaterally in the BLA (2.4 AP, 4.9 ML), with volume of 2µl injected on two levels: 1µl at 6.5 mm and 1µl at 7.5 mm dorsoventral to the dura. Subsequently an optical fiber (200 μm core diameter, Thorlabs, Inc.) was chronically inserted (2.4 AP, 4.9 ML, 6.5 DV). Simultaneous optical stimulation and extracellular recording from CA1 were performed in freely-behaving rats 3 weeks after the surgery. The light power was controlled to be 10-15 mW at the fiber tip. Square-wave pulses of 5 ms were delivered at frequency of 50Hz.

### Spiking ratios

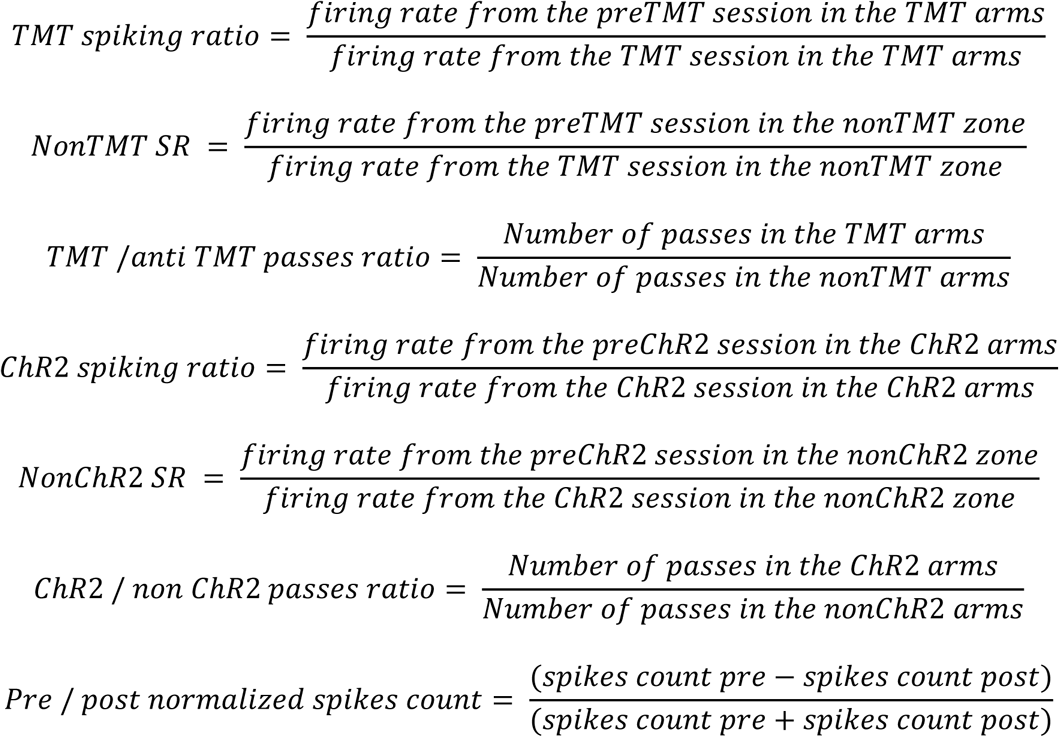

### Center of Mass

The center of mass (COM) was calculated by taking the *x* and *y* averages for the rows and columns of the rate map weighted for firing rate. For the unidirectional place field analyses COM included only the spikes located in the place field defined by the field area smaller than 10% of the maximum firing rate, while for the bidirectional analyses we used all spikes with 20% cut-off. The spatial position of the place cell was defined for all recorded spikes as the center of mass of the firing rate distribution within the maze coordinates. The center of mass of the place cells’ spike distribution is calculated as follows:

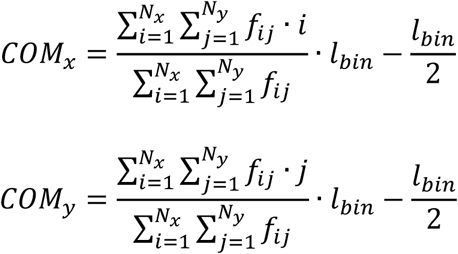

Where *N*_x_, *N*_y_ define the number of bins in the arena in X-, Y-direction; *f*_i,j_ is firing frequency in bin *i, j*; *l*_bin_ is the bin-size.

Given the origin O (O_x_,O_y_), which denotes the northwest (SE) corner of the Cartesian coordinate system, and the direction of the symmetry axis D (D_x_,D_y_), which denotes the line between the southwest (SW) and northeast (NE) corners, the distance of the COM (COM_x_,COM_y_) to the symmetry axis is calculated as follows:

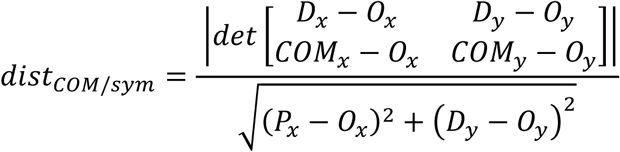

Where *P* is the shortest distance between the COM and the symmetry line. For the rectangular-shaped linear track, the arena borders are defined as the square surrounding all motion tracking sample points with an equal distance to the real limits of the arena at all sides.

Using *dist*_COM_ we calculate the distance between O and P as follows:

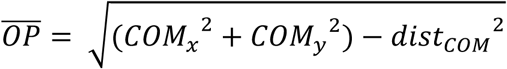

The COM distance normalized by the arena width perpendicular to the symmetry axis through the COM is calculated as:

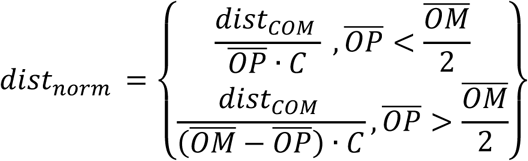

Where 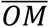 is the diagonal of a square enclosing all motion tracking data points, and *C* is a motion tracking data factor and in this case set to 0.85 for the linear rectangular track.

### Center of Mass Angle

The Center of Mass Angle (COMa) computes the shift of place field COM towards the TMT arms using radial direction in degrees where the axis between the feeding zones denotes 45 degrees. COMa is calculated as follows:

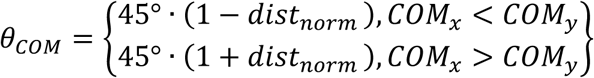

Where *θ*_*COM*_ is center of mass angle; COM_x_, COM_y_ : X-, Y- coordinate of COM, *distnorm* normalized COM distance. The shift in the center of mass (ΔCOM) is the absolute difference between the pre- and post-TMT(ChR2) recordings:

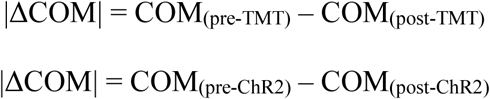

All algorithms were implemented in MATLAB.

### Histology

At the end of the study, brains were removed for histological verification of electrode localization. Rats were deeply anesthetized with sodium pentobarbital (390 mg/kg) and perfused transcardially with ice-cold 0.9% saline followed by 4% paraformaldehyde. Brains were removed, post-fixed in paraformaldehyde for up to 24 hours and cryoprotected in 25% sucrose for >48 hours. Brains were sectioned coronally at 40µm on a freezing microtome. Primary antibody incubations were performed overnight at 4°C in PBS with BSA and Triton X-100 (each 0.2%). The concentration for primary antibodies was anti-CamKIIα 1:500 (Millipore, # 05-532). Sections were then washed and incubated in PBS for 10 minutes and secondary antibodies were added (1:500) conjugated to Alexa Fluor 594 dye (Invitrogen, # A11032) for 2 hours at room temperature.

For visualization, the sections were mounted onto microscope slides in phosphate-buffered water and cover-slipped with Vectashield mounting medium. The YFP fluorescence was evaluated within a selected region that was placed below the fiber tip in an area of 1.5mm x 1.5mm. Fluorescence was quantified based on the average pixel intensity within the selected region (Witten et al., 2011). The stained sections were examined with an Olympus IX81 confocal microscope at 594nm for Alexa Fluor secondary antibody and 488nm for ChR2-YFP. CamKIIα-positive neurons were identified based on the expression of red fluorescence, whereas ChR2-positive neurons were identified by the expression of green fluorescence. Co-localization of Alexa Fluor 594 and YFP was determined manually using ImageJ software.

### Statistical Analysis

Two different approaches were used to calculate the sample size (Karalis et al., 2016). We performed power analyses to establish the required number of animals for experiments in which we had sufficient data on response variables. For experiments in which the outcome of the intervention could not be predetermined, we employed a sequential stopping rule. This approach allows null-hypothesis tests to be used subsequently by analyzing the data at different experimental stages using *t*-tests against Type II error. The experiment was initiated with four animals per group; if *P* < 0.05, the testing was continued with two or three more animals to increase statistical power. In the case of *P* > 0.36, the experiment was discontinued and the null hypothesis was accepted (Karalis et al., 2016). All data were analyzed using SPSS Software. Statistical significance was estimated by using a two-tailed independent samples *t*-test for non-paired data or a paired samples Student *t*-test for paired data. Repeated measures were evaluated with two-way analysis of variance (ANOVA) paired with *post hoc* Bonferroni test. Correlations between data sets were determined using Pearson’s correlation coefficient. The probability level interpreted as significant was fixed at *p* < 0.05. Data are plotted as mean±sem. Dataset of all experimental files is available at https://doi.org/10.6084/m9.figshare.5336026.v1

## Results

### Increase of hippocampal beta amplitude parallels TMT-mediated aversion episodes

To evoke an aversion episode for rats, chronically implanted with tetrodes in hippocampal CA1 region, we used 10% trimethylthiazoline (TMT) applied to a restricted section (10×15 cm^2^) of a rectangular-shaped linear track (Fig. 1A). TMT, a constituent of fox urine, is an innately aversive odor to rodents (Myers and Rinaman, 2005; Kobayakawa et al., 2007). The experimental design consisted of three recording sessions (12 min each), conducted in the range of 48 hours: baseline recording (pre-TMT), recording with TMT located of one of the arms of the track (TMT) and subsequent recording without TMT (post-TMT). During the TMT session the animals (n = 12 rats) avoided to navigate across the TMT-scented section of the track (Fig. 1B, Supplementary Movie 1), resulting in place avoidance (Fig. 1C, paired t-test, n = 12 rats, *t*(11) = 10.4, *P* < 0.001). The experimental design of TMT aversion was designed to evoke in the post-TMT recording session only brief aversive response and subsequent extinction to avoid navigation undersampling in the TMT arms and to identify reliably the place cells’ firing properties. The path sampling data of the pre- and post-TMT sessions show sufficient dwell time spent in both TMT- and non-TMT arms with sufficient path sampling for all rats (Fig. 1C, Table 1). The association of the TMT arms with the aversive odor was evident during the first minute of navigation in the post-TMT session (Fig. 1D, paired t-test, n = 12, *t*(11) = −4.244, *P*= 0.001). The average number of passes (2.58 ± 0.3) was significantly lower for the TMT arms compared to the non-TMT arms (5.16 ± 0.7). To identify if the olfaction-mediated aversion was restricted to the TMT section of the maze or if it was occurring over the entire recording arena we analyzed the amplitude of hippocampal beta frequency band (15-40Hz, Fig. 1E, Fig. 1F) for each arm of the track (Fig. 1G). Successful odor discrimination is identified by hippocampal oscillations in beta frequency range (Igarashi et al., 2014) and therefore, the increase of beta amplitude is a reliable indicator of aversive odors processing by the hippocampal network. The hippocampal beta amplitude (Fig. 1H, one-way ANOVA, n = 12, F_(1,7)_ = 3.1, *P* = 0.005) and frequency (Fig. 1I, one-way ANOVA, n = 12, F_(1,7)_ = 3.8, *P* = 0.002) expressed dependence on the animal’s whole body linear speed. Concurrently, the TMT-induced aversion impacted the animals’ linear speed (Supplementary Movie 1). To avoid the bias of speed on beta parameters we analyzed the effect of TMT on beta oscillations for different speed bands. The highest increase of beta amplitude was evident during the passes on the arm with TMT odor (TMT arm 1, Fig. 1J, two-way ANOVA with Bonferroni post-hoc test, between groups, n = 12, F_(2,10)_ = 17.5, *P* < 0.001) with significant increase for five speed ranges. Significant beta amplitude increase was present also in the second arm between the food zones, adjacent to the TMT arm (TMT arm 2, Fig. 1K, two-way ANOVA with Bonferroni post-hoc test, between groups, F_(2,10)_ = 24.3, *P* < 0.001). The opposing loop between the food zones including non-TMT arm 1 and non-TMT arm 2 was characterized with non-significant changes of beta amplitude (Fig. 1L,M, two-way ANOVA with Bonferroni post-hoc test, between groups, n = 12, for non-TMT arm 1: F_(2,10)_ = 3.0, *P* = 0.094 and for non-TMT arm 2: F_(2,10)_ = 1.3, *P* = 0.295). We found no significant effect of TMT on the frequency of beta oscillation (Fig. 1N, O, TMT arm 1: two-way ANOVA, between groups, with Bonferroni post-hoc correction, n = 12, F_(2,10)_ = 0.7, *P* = 0.493; TMT arm 2: n = 12, F_(2,10)_ = 1.2, *P* = 0.349; non-TMT arm 1: n = 12, F_(2,10)_ = 2.1, *P* = 0.164; non-TMT arm 2: n = 12, F_(2,10)_ = 1.8, *P* = 0.213). These data indicate that the effect of TMT on the hippocampal local field activity was restricted to two arms of the rectangular-shaped linear track (TMT arms).

**Figure 1.**
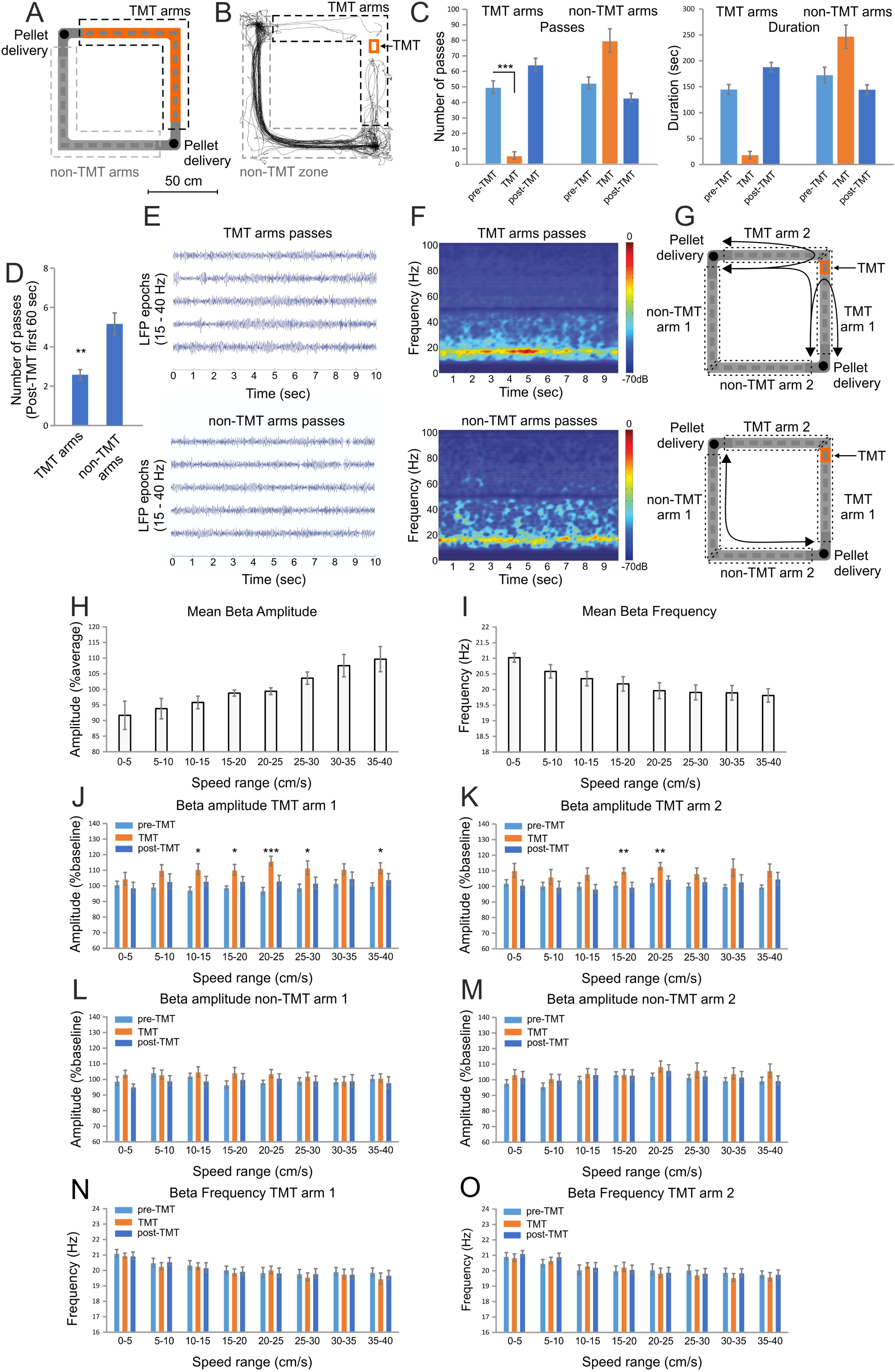
Hippocampal beta amplitude increase during odor-triggered place avoidance. ***A***, Behavioral set-up of rectangular-shaped linear track. Locations of the TMT are marked with red in the north and east arms of the track. The black dashed line indicates the TMT arms, while grey dashed line marks the non-TMT arms. To match the TMT protocol with the subsequent ChR2 photostimulation protocol we applied the TMT scent papers to all locations across the TMT arms. ***B***, Navigation path of sample animal during the TMT session. Note the place avoidance of the TMT arms. The grey dashed line marks the non-TMT zone. ***C***, Number of passes (left graph) and duration in seconds (right graph) counted in the TMT arms vs non-TMT arms during the pre-TMT-, TMT- and post-TMT sessions, \*\*\**P* < 0.001. Error bars, mean ± s.e.m. ***D***, Number of passes through the TMT- and non-TMT arms for the first 60 seconds of the post-TMT session. \*\**P* < 0.01. ***E***, Representative band pass filtered (15 – 40Hz) local field potential (LFP) recorded during the passes in the TMT arms (top panel) and in the non-TMT arms (bottom panel). Time 0 indicates the onset of the path trajectory starting from pellet delivery location. ***F***, Representative averaged band pass filtered (15 – 40Hz) color-coded power spectrogram of all passes in the TMT arms (top panel) and in the non-TMT arms (bottom panel). Time 0 indicates the onset of the path trajectory starting from pellet delivery location. The averaged power-spectrogram includes the variability of passes duration and speed during the navigation across the arms. ***G***, Spatial dissociation of odor perception for in TMT arms (top panel) and in the non-TMT arms (bottom panel). The arm with TMT scent paper (marked with red) was named TMT arm 1. The adjacent arm, part of the same food-navigation loop, was TMT arm 2. The opposite of the TMT arm 1 was named non-TMT arm 1, while the opposite of the TMT arm 2 was non-TMT arm 2. The scent was detected by the animal at different locations during the navigation passes across the TMT arms. Black arrows indicate the possible path trajectories of the animals. ***H***, Mean beta amplitude and ***I***, mean beta frequency measured during different whole body speed ranges of 0 – 40 cm/s in bins of 5 cm/s. ***J***, Beta amplitude for TMT arm 1 and ***K*** TMT arm 2 (right) as percent of the pre-TMT session values for the entire track. \*\*\**P* < 0.001, \*\**P* < 0.01, \**P* < 0.05 Error bars, mean ± s.e.m. ***L***, Beta amplitude for non-TMT arm 1 and ***M***, non-TMT arm 2 as percent of the pre-TMT session values for the entire track. The amplitude values are presented as a function of the animal’s whole body speed, where beta amplitude is evaluated for speed range of 0 – 40 cm/s in bins of 5 cm/s. Error bars, mean ± s.e.m. ***N***, Beta frequency for TMT arm 1 and ***O***, TMT arm 2 for speed range of 0 – 40 cm/s in bins of 5 cm/s. Error bars, mean ± s.e.m.

**Table 1:** Path sampling from recordings with aversive experimental design. Number of passes and dwell time (duration in seconds) measured in the south and west (SW), and north and east (NE) arms for TMT- and ChR2- groups of animals during the pre-TMT(ChR2), TMT(ChR2)- and post-TMT(ChR2) recording sessions. The rats exposed to TMT include TMT-NE and TMT-SW-groups.

| TMT-NE<br>Rat # | Pre-TMT session |  |  |  | TMT-NE session |  |  |  | Post-TMT session |  |  |  |
| --- | --- | --- | --- | --- | --- | --- | --- | --- | --- | --- | --- | --- |
|  | Passes |  | Duration (sec) |  | Passes |  | Duration (sec) |  | Passes |  | Duration (sec) |  |
|  | SW | NE | SW | NE | SW | NE | SW | NE | SW | NE | SW | NE |
| Rat 1 | 38 | 42 | 102.2 | 112.7 | 50 | 0 | 150.2 | 0 | 42 | 31 | 134.6 | 107.5 |
| Rat 2 | 33 | 41 | 156.9 | 181.4 | 91 | 9 | 266.4 | 65.4 | 60 | 51 | 187.5 | 160.7 |
| Rat 3 | 31 | 48 | 122.8 | 218.8 | 62 | 0 | 229.4 | 0 | 74 | 24 | 241.0 | 91.92 |
| Rat 4 | 53 | 59 | 153.1 | 183.1 | 70 | 0 | 272.2 | 0 | 47 | 43 | 189.5 | 154.9 |
| Rat 5 | 48 | 54 | 150.4 | 194.3 | 78 | 0 | 260.7 | 0 | 75 | 37 | 233.2 | 116.5 |
| Rat 6 | 64 | 80 | 163.1 | 210.9 | 101 | 0 | 308.8 | 0 | 72 | 60 | 194.2 | 189.9 |
| TMT-SW<br>Rat # | Pre-TMT session |  |  |  | TMT-SW session |  |  |  | Post-TMT session |  |  |  |
|  | Passes |  | Duration (sec) |  | Passes |  | Duration (sec) |  | Passes |  | Duration (sec) |  |
|  | SW | NE | SW | NE | SW | NE | SW | NE | SW | NE | SW | NE |
| Rat 7 | 31 | 50 | 105.8 | 157.9 | 15 | 47 | 50.9 | 187.9 | 38 | 42 | 112.0 | 153.0 |
| Rat 8 | 56 | 59 | 170.8 | 152.5 | 0 | 95 | 0 | 350.3 | 41 | 63 | 136.5 | 166.9 |
| Rat 9 | 40 | 39 | 160.7 | 107.1 | 0 | 106 | 0 | 258.7 | 49 | 87 | 152.6 | 159.4 |
| Rat 10 | 61 | 51 | 167.7 | 132.7 | 16 | 46 | 30.2 | 79.2 | 44 | 73 | 154.2 | 180.6 |
| Rat 11 | 60 | 52 | 175.5 | 138.4 | 24 | 82 | 52.7 | 248.3 | 42 | 65 | 163.2 | 188.4 |
| Rat 12 | 59 | 80 | 186.1 | 198.6 | 5 | 130 | 17.1 | 346.1 | 55 | 72 | 190.9 | 224 |
| ChR2-SW<br>Rat # | Pre-ChR2 session |  |  |  | ChR2-SW session |  |  |  | Post- ChR2 session |  |  |  |
|  | Passes |  | Duration (sec) |  | Passes |  | Duration (sec) |  | Passes |  | Duration (sec) |  |
|  | SW | NE | SW | NE | SW | NE | SW | NE | SW | NE | SW | NE |
| Rat 1 | 46 | 34 | 140.7 | 105.2 | 27 | 49 | 175.4 | 119.5 | 29 | 13 | 88.0 | 181.2 |
| Rat 2 | 31 | 54 | 98.4 | 153.9 | 23 | 52 | 75.4 | 159.8 | 36 | 54 | 166.1 | 155.4 |
| Rat 3 | 26 | 46 | 84.7 | 102.2 | 21 | 41 | 84.3 | 160.6 | 31 | 54 | 107.8 | 167.1 |
| Rat 4 | 35 | 41 | 118.3 | 131.0 | 11 | 38 | 146.1 | 167.0 | 45 | 47 | 170.1 | 135.8 |
| Rat 5 | 33 | 48 | 100.3 | 130.7 | 15 | 38 | 101.6 | 128.4 | 47 | 44 | 113.6 | 96.0 |
| Rat 6 | 36 | 42 | 114.6 | 130.5 | 13 | 40 | 201.7 | 197.4 | 38 | 40 | 169.7 | 151.5 |

### The place cell’s activity in the TMT arms during aversion episodes correlates to the degree of field reconfiguration

We next explored if the exposure to TMT increased remapping propensity of the place cells, measured by the shift of the center of mass (ΔCOM) for all spikes. ΔCOM is an efficient approach for field plasticity evaluation because this parameter represents spiking as a function of the occupancy for each pixel and detects both spatial and rate remapping (Knierim, 2002; Lee et al., 2004). The average ΔCOM between the pre-TMT and post-TMT recordings for the place cells from the TMT group was 12.44 ± 0.8 cm (Fig. 2A). This was a significant increase compared to the ΔCOM, examined from animals undergoing control recordings with indifferent odor: 10% ethanol, with baseline value of 6.32 ± 0.6 cm (unpaired t-test, control group n = 57, TMT group n = 106, *t*(161) = 5.2, *P* < 0.001). Furthermore, the ratio of the TMT-over non-TMT number of passes for the first 60 seconds significantly correlated to the degree of ΔCOM (Fig. 2B, Pearson’s r = −0.656, *P* = 0.020, n = 12). We investigated if the intra- and extra-field spiking activity of individual place cells during the TMT episodes (Fig. 2C) relates to subsequent shift of the place cell’s center of mass angle (ΔCOMa). This parameter estimates the proximity of the center of mass to the TMT zone across the main axis of the track (see Methods & Materials), between 0° for SW corner and 90° for NE corner. The TMT session induced variable ΔCOMa between the pre-TMT and post-TMT sessions for the NE group of animals (Fig. 2D) as well as for the SW group of animals (Fig. 2E). We correlated the degree of ΔCOMa to the change of the firing rate of the place cells within the TMT arms. The change of the firing rate was calculated by the ratio of the baseline over the TMT session firing rate from the recorded spikes. We found significant negative correlation between ΔCOMa and the TMT spiking ratio of the mean firing rate measured for the TMT arms (Fig. 2F, Pearson’s r = −0.276, *P* = 0.004, n = 106). Concurrently, no significance was evident for the correlation between ΔCOMa and the non-TMT mean spiking ratio measured from the non-TMT zone (Fig. 2G, Pearson’s r = −0.016, p = 0.871, n = 107). Similarly, the correlation between ΔCOMa and the spiking ratio of the peak firing rate was significant for the TMT arms (Fig. 2H, Pearson’s r = −0.271, *P* = 0.005, n = 106), but not for the non-TMT zone (Fig. 2I, Pearson’s r = −0.083, *P* = 0.394, n = 107). In the control group of rats (Fig. 3A,B) we found no significant correlation between ΔCOMa (Fig. 3C,D) and the mean spiking ratio for the ethanol arms (Fig. 3G, Pearson’s r = −0.001, *P* = 0.997, n = 31) and for the non-ethanol zone (Fig. 3H, Pearson’s r = - 0.041, *P* = 0.822, n = 32). The correlations between ΔCOMa and peak spiking ratios were also non-significant (3I, Pearson’s r = −0.092, *P* = 0.621, n = 31; Fig. 3J, Pearson’s r = −0.062, *P* = 0.734, n = 32). These data demonstrate that the cells’ spiking during aversive episodes is related to subsequent place field remapping.

**Figure 2.**
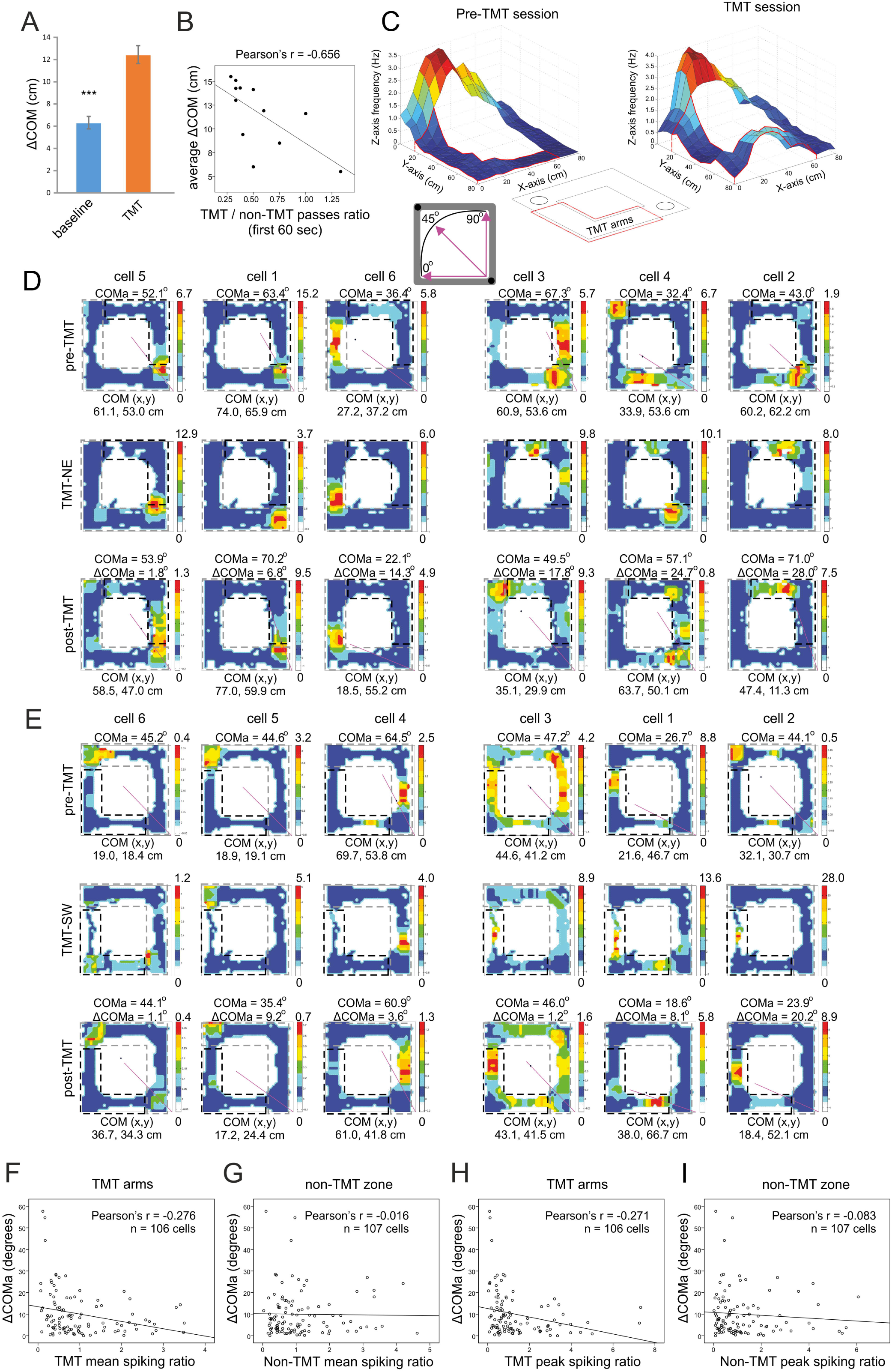
Individual place field reconfiguration after exposure to TMT. ***A***, Center of mass shift (ΔCOM) of place cells recorded from baseline group of animals exposed to familiar odor (10% ethanol) and group of rats exposed to innately aversive odor (10% TMT). \*\*\**P* < 0.001. Error bars, mean ± s.e.m. ***B***, Correlation between the average ΔCOM for each rat and the ratio of the TMT-over the non-TMT arms passes for the first 60 seconds of the post-TMT session. ***C***, Left: Color-coded 3D-spatial map of a sample place cell with place field located outside the TMT arms (the TMT arms are marked with red line), recorded during pre-TMT session. X- and Y-axes represent the coordinates of the recording arena, while Z-axis represents the firing rate of the recorded neuron. Blue colors represent low-, while red colors represent high firing rate. Right: Color-coded 3D-spatial map of a sample place cell with place field located outside the TMT arms (the TMT arms are marked with red line), recorded during TMT session. The inset below shows with red line the position of the TMT arms (circles indicate the pellet delivery corners). The spiking TMT ratio represents the mean (peak) firing rate of the pre-TMT session over the TMT session. The inset below shows the radial representation of COMa. The straight purple line starting from the southeast corner of the track and it indicates the COM position in respect to the main axis of the track positioned between the food zones (marked with black dots, in the SE and NW corners). Value of 45° indicates COM evenly distributed across the main axis of the track, 0° indicates COM fully distributed within the SW section and 90° - in the NE section of the track. ***D***, Six sample place cells recorded from animal of the NE group during pre-TMT session (upper panels), TMT session (middle panels) and post-TMT session (lower panels). The straight purple line denotes the center of mass angle (COMa) for each cell between SW at 0° and NE at 90°. Note that some of the place cells exhibited higher firing rate in the TMT arms during the TMT sessions (panels positioned on the right half) compared to place cells with little or no spiking (panels positioned on the left half). ***E***, Six sample place cells recorded from animal of the SW group during pre-TMT session (upper panels), TMT session (middle panels) and post-TMT session (lower panels). ***F***, Correlation between ΔCOMa and TMT spiking ratio based on the mean firing rate of the place cells’ spikes in the TMT arms. ***G***, Correlation between ΔCOMa and non-TMT spiking ratio based on the mean firing rate of the place cells’ spikes in the non-TMT zone. ***H***, Correlation between ΔCOMa and the spiking ratio based on the peak firing rate for the spikes in the TMT arms. ***I***, Correlation between ΔCOMa and the spiking ratio based on the peak firing rate for the spikes in the non-TMT zone.

**Figure 3.**
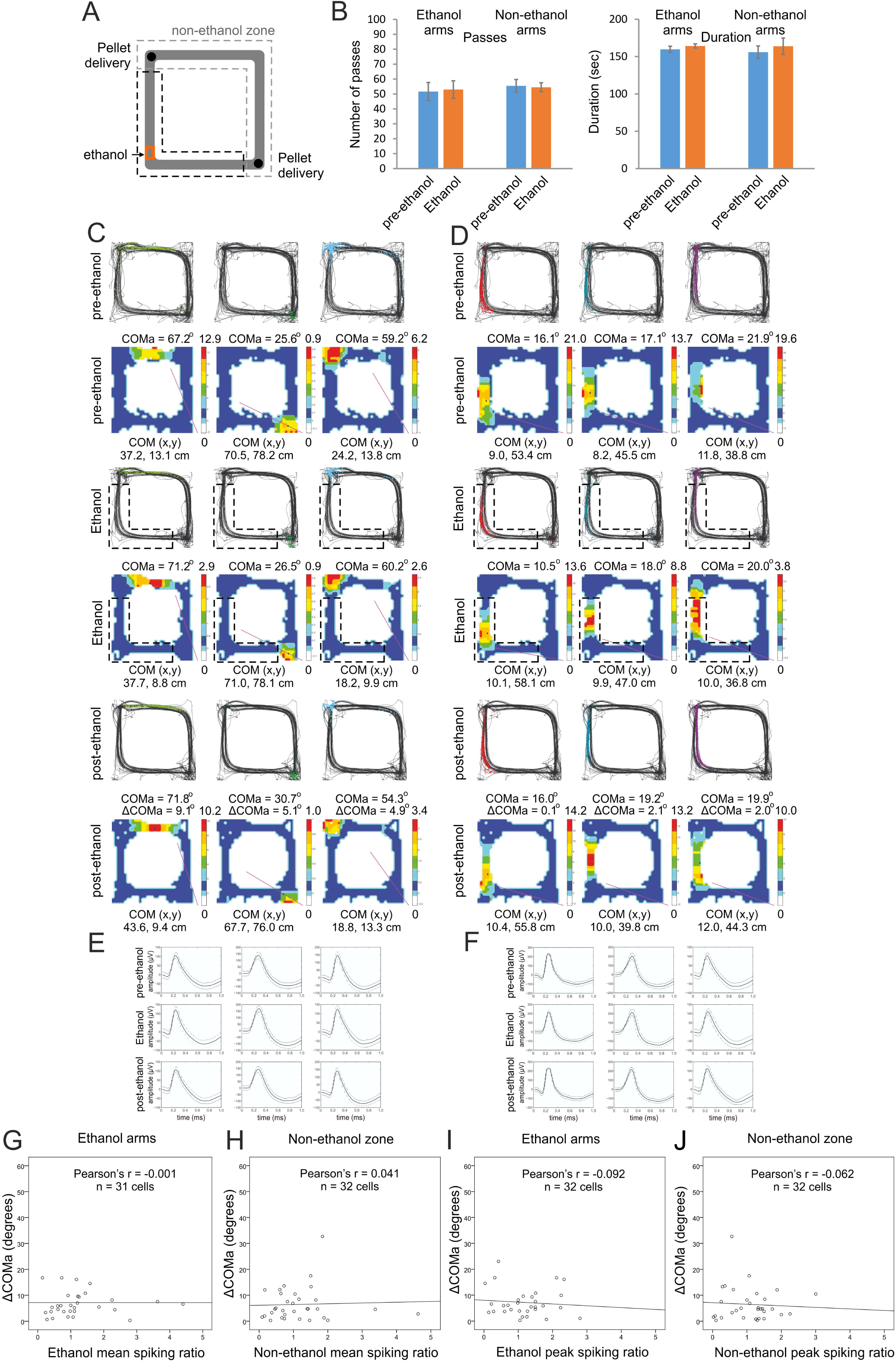
Place field stability in control conditions. ***A***, Schematic representation of the application of a familiar scent, ethanol (marked with red) in the SW arms of rectangular-shaped linear track for the control group of animals. ***B***, Number of passes (left panel) and duration in seconds (right panel) counted in the ethanol arms (left) and non-ethanol arms (right) for the control group of rats, before (blue) and during (red) the odor exposure. Error bars, mean ± s.e.m. ***C***, Three place fields from sample animal with fields located outside the ethanol arms during pre- (top), ethanol (middle) and post-ethanol sessions (bottom). For each session the upper panels show the animal trajectory with spikes, marked with colored dots, while the lower panels show color-coded firing rate. ***D***, Three place fields from same animal with fields located inside the ethanol arms during pre- (top), ethanol (middle) and post-ethanol sessions (bottom). For each session the upper panels show the animal trajectory with spikes, marked with colored dots, while the lower panels show color-coded firing rate. ***E***, Waveforms of the place cells shown in C, and ***F***, waveforms of the cells shown in D, respectively. The solid line shows the average waveform shape; the dashed lines show the 1 SD confidence intervals. ***G***, Correlation between ΔCOMa and the spiking ratio based on the mean firing rate of the place cells’ spikes in the ethanol arms. ***H***, Correlation between ΔCOMa and the spiking ratio based on the mean firing rate of the place cells’ spikes in the non-ethanol zone. ***I***, Correlation between ΔCOMa and the spiking ratio based on the peak firing rate for the spikes in the ethanol arms. ***J***, Correlation between ΔCOMa and the spiking ratio based on the peak firing rate for the spikes in the non-ethanol zone.

### The extra-field spiking during aversion episodes determines the degree of field plasticity

We compared the field reconfiguration of the place cells with fields located in the TMT arms (with spiking ratio including intra-field spikes, Fig. 4A) to the remapping of the cells with place fields outside the TMT arms (with spiking ratio including only extra-field spikes, Fig. 4B). The place field was defined as the region of the arena in which the firing rate of the place cell was > 20% of the peak firing frequency (Brun et al., 2002). The extra-field TMT spiking ratio highly correlated to ΔCOMa for both the mean (Fig. 4E, Pearson’s r = −0.465, *P* < 0.001, n = 54) and for the peak firing rate (Fig. 4I, Pearson’s r = −0.453, *P* < 0.001, n = 54). However, the intra-field TMT spiking ratio showed no significant correlation to ΔCOMa for the mean firing rate (Fig. 4F, Pearson’s r = 0.013, *P* = 0.925, n = 52) as well as for the peak firing rate (Fig. 4J, Pearson’s r = −0.055, *P* = 0.698, n = 52). The dissociation of the spikes into TMT extra- and intra-field groups had no effect on the place cells activity outside the TMT arms during the TMT sessions. No significant correlation was evident for the mean and peak firing rate of the intra- (Fig. 4G, Pearson’s r = −0.024, *P* = 0.864, n = 55; Fig. 4K, Pearson’s r = −0.119, *P* = 0.387, n = 55) and extra-field spikes (Fig. 4H, Pearson’s r = 0.027, *P* = 0.852, n = 52; Fig. 4L, Pearson’s r = 0.062, *P* = 0.661, n = 52) in the non-TMT zone. These results show that the degree of increase in extra-field spiking during the TMT sessions predicts the degree of place field plasticity. Furthermore, the degree of increase extra-field spiking of these place cells during the TMT sessions predicted the degree of place field remapping.

**Figure 4.**
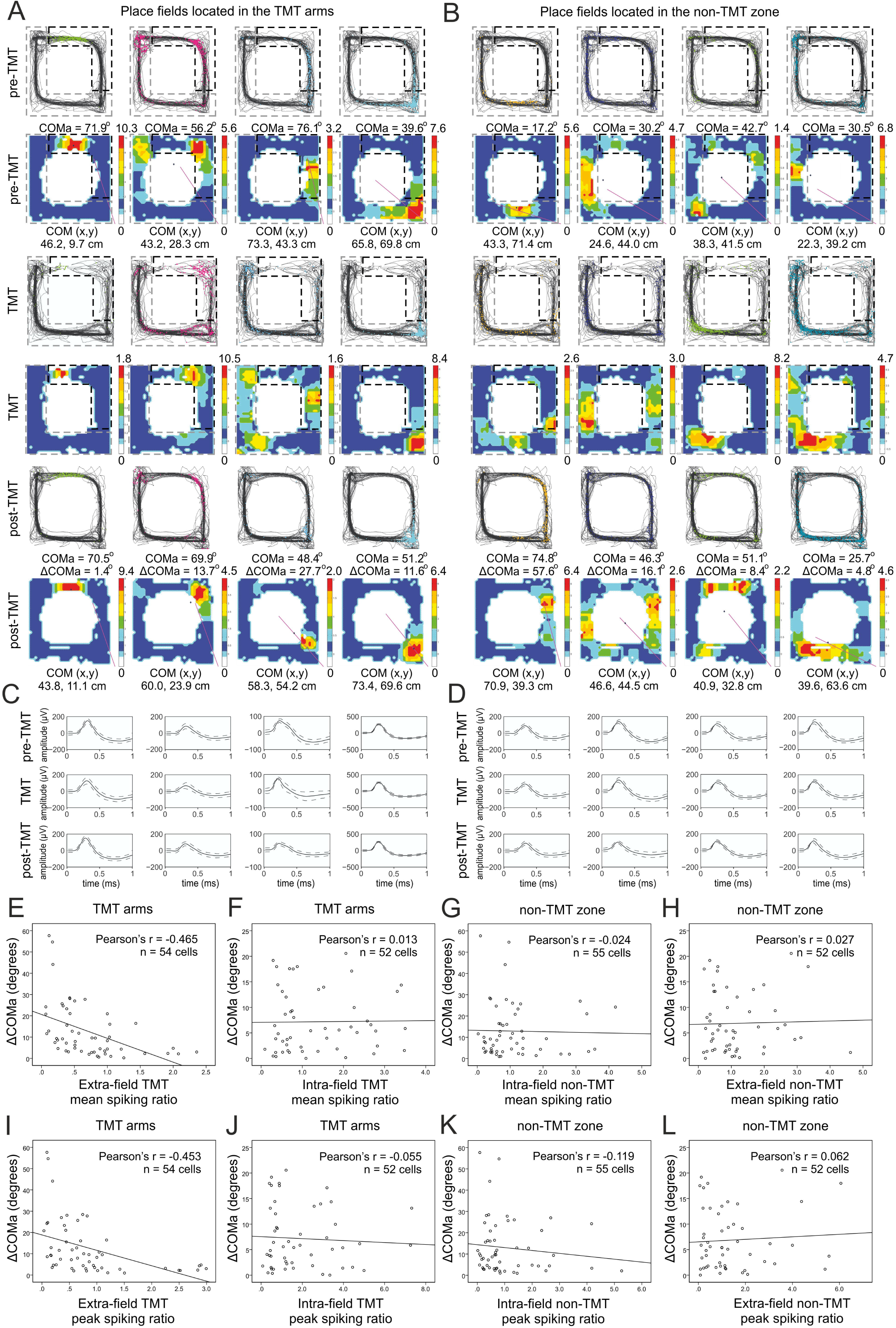
Increased extra-field place cell spiking during TMT exposure predicts spatial field reconfiguration. ***A***, Four place fields from sample animal with fields located in the TMT arms during pre-TMT sessions (top), TMT (middle) and post-TMT sessions (bottom). For each session the upper panels show the animal trajectory with spikes, marked with colored dots, while the lower panels show color-coded firing rate. ***B***, Four place fields from the same animal with fields outside the TMT arms. ***C***, Waveforms of the place cells from the pre-TMT, TMT and post-TMT sessions shown in A, and ***D***, waveforms of the cells shown B, respectively. The solid line shows the average waveform shape; the dashed lines show the 1 SD confidence intervals. ***E***, Correlation between ΔCOMa and TMT mean spiking ratio for the place cells located outside the TMT arms (extra-field spikes), Supplementary Table 1. *F*, Correlation between ΔCOMa and TMT mean spiking ratio for the place cells located inside the TMT arms (intra-field spikes), Supplementary Table 2. ***G***, Correlation between ΔCOMa and non-TMT mean spiking ratio for the place cells located inside the non-TMT arms (intra-field spikes), Supplementary Table 3. ***H***, Correlation between ΔCOMa and non-TMT mean spiking ratio for the place cells located outside the non-TMT zone (extra-field spikes), Supplementary Table 4. ***I***, Correlation between ΔCOMa and TMT peak spiking ratio for the place cells located outside the TMT arms (extra-field spikes), Supplementary Table 1. ***J***, Correlation between ΔCOMa and TMT peak spiking ratio for the place cells located inside the TMT arms (intra-field spikes), Supplementary Table 2. ***K***, Correlation between ΔCOMa and non-TMT peak spiking ratio for the place cells located inside the non-TMT zone (intra-field spikes), Supplementary Table 3. ***L***, Correlation between ΔCOMa and non-TMT peak spiking ratio the place cells located outside the non-TMT zone (extra-field spikes), Supplementary Table 4.

### The center of mass shift occurs in both directions of navigation along the track

We next aimed to precisely identify the degree of place field COM shift between recording sessions. The angle calculation of COM in 2D space is prone to higher variability compared to the one-directional COM calculation since the distance covered per unit angle varies as a function of the radial distance. For this purpose we computed the degree of field remapping and firing rate remapping after the path of the animals was linearized (Fig. 5). Place fields on linear tracks display differential firing depending on the direction of the animal’s movement (McNaughton et al., 1983). Thus, we also separated the place fields into clockwise and counter-clockwise trajectories (Fig. 5A,C,E Fig. 5B,D,F). For the unidirectional place fields we used an alternative place field cut-off, which includes the field bins with firing rate greater than 10% of the maximum firing rate (see Methods & Materials). We compared the spiking parameters of the place fields between: 1) pre-ethanol session (Fig. 5A,B left panels) and post-ethanol session (Fig. 5A,B right panels); between 2) pre-TMT session for the SW group (Fig. 5C,D left panels) and post-TMT session (Fig. 5C,D right panels), and between 3) pre-TMT session for the NE group (Fig. 5E,F left panels) and post-TMT session (Fig. 5E,F right panels). There was no significant difference in the change of the mean firing rate between the ethanol, TMT-NE and TMT-SW groups for the clockwise (Fig. 5G, one-way ANOVA with Bonferroni post-hoc correction, n = 6, F_(2,128)_ = 0.257, *P* = 0.773) and counter-clockwise groups (Fig. 5J, one-way ANOVA with Bonferroni post-hoc correction, n = 6, F_(2,118)_ = 0.164, *P* = 0.849). Similarly, the change of the peak firing rate between the three groups was not significant for the clockwise (Fig. 5H, one-way ANOVA with Bonferroni post-hoc correction, n = 6, F_(2,128)_ = 0.886, *P* = 0.886) or for counter-clockwise fields (Fig. 5K, one-way ANOVA with Bonferroni post-hoc correction, n = 6, F_(2,118)_ = 0.118, *P* = 0.889). The linearized COM shifted on average with 5.10 ± 0.5 cm for the clockwise- (Fig. 5I) and 5.37 ± 0.5 cm for the counter-clockwise fields (Fig. 5L) in the ethanol-exposed group. Exposure to TMT evoked ΔCOM of 24.24 ± 4.6 cm for clockwise- and 24.47 ± 4.4 cm for counter-clockwise fields for the TMT-NE group, and 21.84 ± 3.3 cm for clockwise- (Fig. 5I) and 24.65 ± 4.0 cm for counter-clockwise fields (Fig. 5L) for the TMT-SW group. This shift was significantly higher compared to the control group for clockwise- (one-way ANOVA with Bonferroni post-hoc correction, n = 6, F_(2,128)_ = 11.52, *P* < 0.001) and counter-clockwise fields (one-way ANOVA with Bonferroni post-hoc correction, n = 6, F_(2,118)_ = 10.105, *P* < 0.001). These data show that TMT evokes potent field remapping and negligible rate remapping for both directions of navigation.

**Figure 5.**
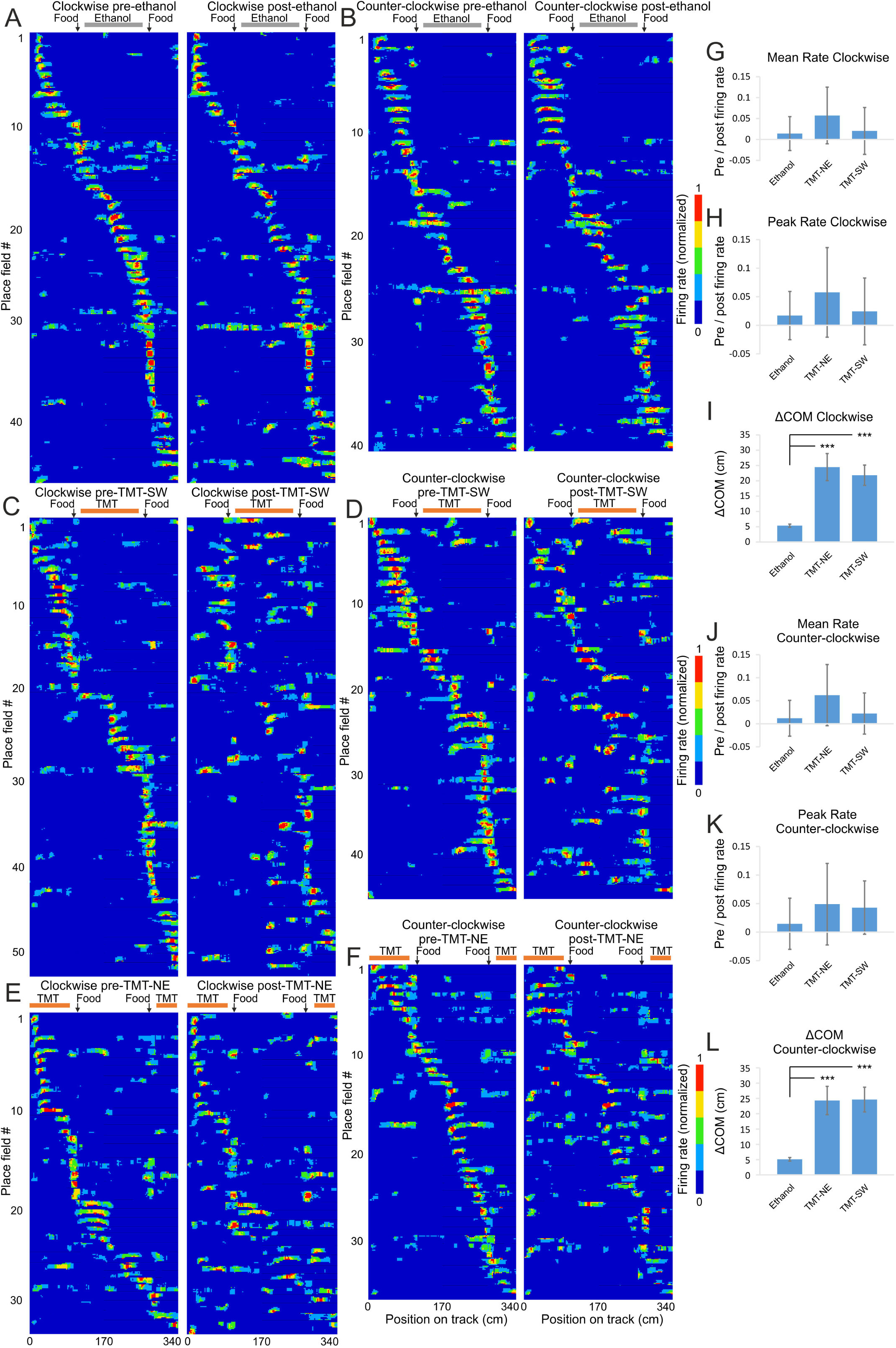
Field but not rate remapping after aversive experience. ***A***, Color-coded linearized map showing location of CA1 place fields before (left panel) and after (right panel) exposure to ethanol for clockwise, Supplementary Table 5 and ***B***, counter-clockwise direction of movement, Supplementary Table 6. Each line shows activity of one place cell (86 datasets in total from 63 place cells). The horizontal grey bar indicates the ethanol zone during the exposure session, while black vertical arrows indicate the location of the food delivery. ***C***, Linearized map before (left panel) and after (right panel) exposure to TMT in the SW section of the track for clockwise, Supplementary Table 7 and ***D***, counter-clockwise direction of movement, Supplementary Table 8. Each line shows activity of one place cell (98 datasets in total from 66 place cells). The horizontal red bar indicates the TMT zone during the exposure session, while black vertical arrows indicate the location of the food delivery. ***E***, Linearized maps before (left panel) and after (right panel) exposure to TMT in the NE section of the track for clockwise, Supplementary Table 9 and ***F***, counter-clockwise direction of movement, Supplementary Table 10. Each line shows activity of one place cell (68 datasets in total from 40 place cells). ***G***, Comparison of the place field mean spiking before and after exposure to TMT. The pre/post normalised count represents decrease (positive) and increase (negative values) for the mean firing rate of clockwise fields for ethanol, TMT-NE- and TMT-SW groups. Error bars, mean ± s.e.m. ***H***, Comparison of the place field peak spiking before and after exposure to TMT of clockwise fields for the same groups. Error bars, mean ± s.e.m. ***I***, Center of mass shift (ΔCOM) after exposure to TMT of clockwise fields for ethanol, TMT- NE- and TMT-SW groups. Error bars, mean ± s.e.m. \*\*\**P* < 0.001. ***J***, Comparison of the place field mean spiking before and after exposure to TMT for counter-clockwise fields. Error bars, mean ± s.e.m. ***K***, Comparison of the place field peak spiking before and after exposure to TMT for counter-clockwise fields. Error bars, mean ± s.e.m. ***L***, ΔCOM after exposure to TMT for counter-clockwise fields. Error bars, mean ± s.e.m. \*\*\**P* < 0.001.

### BLA photostimulation mediates aversion-triggered shift of the center of mass

Basolateral amygdala (BLA) activation is known to trigger aversive behavior (Davis, 1992). We investigated if BLA innervation of hippocampal place cells mediates the aversion induced field plasticity and if optogenetic activation BLA excitatory neurons will exert an effect similar to the TMT-evoked pattern of reconfiguration. To exert spatial control of the BLA neuronal activity we injected a viral construct AAV-CaMKIIα-hChR2-YFP in the BLA of Lister-Hooded rats (Fig. 6A). Delivery of blue light (473 nm) excited the spiking of neurons infected with AAV-CaMKIIα-hChR2-YFP (Fig. 6B,C) and induced concurrently aversive behavior (Supplementary Movies 2, 3). The majority of the photostimulated BLA neurons were CamKIIα-positive: 87 ± 8% of neurons that expressed yellow fluorescent protein (YFP) also expressed CamKIIα, while 64 ± 5% of neurons that expressed CamKIIα also expressed YFP. We applied optogenetic stimulation (50Hz, trains of 12 pulses, 0.5Hz inter-train interval) in south and west arms of the track (ChR2 arms, Fig. 6D) and observed place avoidance in the ChR2 arms (Fig. 6E, paired t-test, n = 6 rats, *t*(5) = 5.0, *P* = 0.004). The post-ChR2 session recollected the place aversion of the animals in the first minute of the recording (Fig. 6F, paired t-test, n = 6, *t*(5) = −3.796, p = 0.013). BLA photostimulation affected the synchronization of hippocampal local field oscillations across the stimulation trials (Fig. 6G), where the power of the event-related potential increased in the range of 5 – 8 Hz (Fig. 6H). The phase-locking value is a parameter that measures the degree of local field synchrony between all stimulation epochs (see Materials & Methods). The phase-locking values were significantly higher for BLA light pulse delivery (Fig. 6I, 0.35 ± 0.04) compared to the shuffled BLA data (0.09 ± 0.07, paired t-test, n = 120 trials, *t*(119) = 3.0, *P* = 0.005), as well as compared to the control YFP group (0.01 ± 0.06, unpaired t-test, n = 120 trials, *t*(118) = 4.8, *P* = 0.009). We next compared the change of the intra-field and extra-field spiking of place cells that fire in both ChR2-arms and non-ChR2 zone. The spike count was examined in epochs of 100, 250, and 500ms pre- and post-BLA stimulation protocol onset. The positive value of the normalized count (0.03 ± 0.03) showed that the activation of the amygdala afferents resulted in a tendency of decreased intra-field spiking rate in the first 100ms post-BLA photostimulation (Fig. 6J). Concurrently, the extra-field spiking increased (represented by the negative value of the normalized count, −0.11 ± 0.04), which was significantly different from the extra-field normalized count for the first 100ms (paired t-test, n = 74 cells, *t*(73) = 3.2, *P* = 0.002), but not the 250ms (paired t-test, n = 74 cells, *t*(73) = 1.1, *P* = 0.254) and 500ms post-BLA photostimulation (paired t-test, n = 74 cells, *t*(73) = 1.2, *P* = 0.247). We compared the COM angle shift (ΔCOMa) between the pre- and post-ChR2 sessions for the place cells with fields located in the ChR2 arms (with intra-field spikes) and for the place cells with fields located in the non-ChR2 zone (with extra-field spikes only). The average ΔCOMa of 7.57 ± 1.1 from cells with fields located in the ChR2 arms was significantly lower than ΔCOMa of 12.64 ± 1.6 from cells located in the non-ChR2 zone (Fig. 6K, unpaired t-test, extra-field group, n = 38 cells, intra-field group, n = 42 cells, *t*(78) = 5.0, *P* = 0.011). ΔCOMa for the extra-field spiking cells also differed significantly between the ChR2- and YFP control group of rats (unpaired t-test, ChR2 extra-field group, n = 38 cells, YFP extra-field group, n = 32 cells, *t*(68) = 5.3, *P* = 0.010). The application of AAV-CaMKIIα-YFP in control rats (Fig. 7A) evoked no behavioral response (Fig. 7D,E, paired t-test, n = 6 rats, *t*(5) = 1.0, *P* = 0.358), electrophysiological response (Fig. 7B,C,F-H) or remapping (Fig. 8A,B). These data confirm BLA photoactivation as a reliable experimental protocol for behavioral place aversion and place field plasticity.

**Figure 6.**
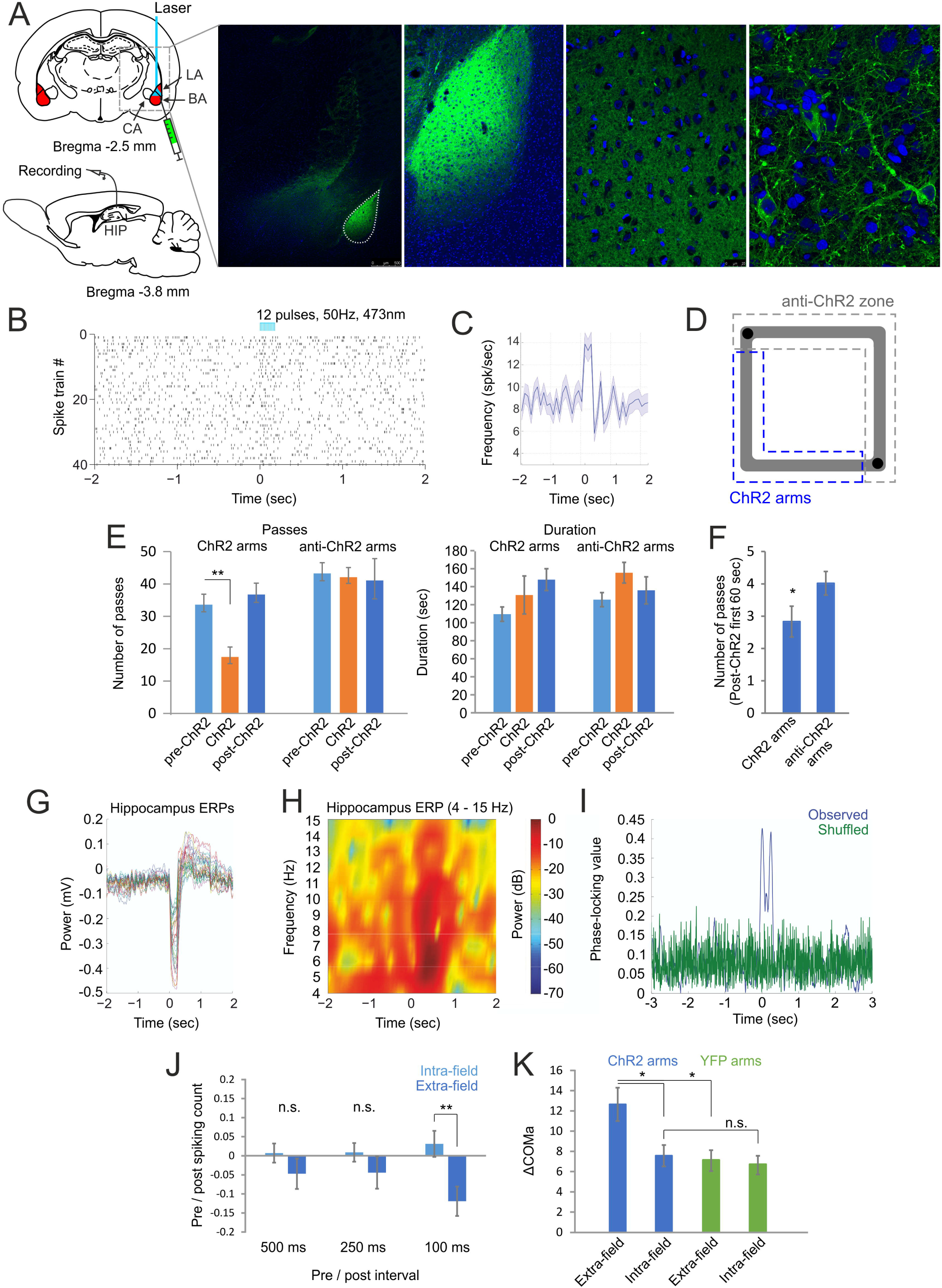
Photostimulation of basolateral complex of amygdala evokes spatial aversion. ***A***, Atlas schematic shows the injection site of AAV-CaMKIIα-hChR2-YFP and the optic fiber location in the basolateral complex of amygdala (BA – basal nucleus, LA – lateral nucleus of amygdala, CA – central nucleus of amygdala). The confocal images on the right show increasing levels of magnification of the YFP-expressing neurons (green) in BLA is histology with DAPI staining (blue). ***B***, Raster plot from 40 repetitions and ***C***, averaged firing frequency of optically evoked time-locked excitation of a BLA cell. Time 0 indicates the delivery of the first train of the stimulation protocol. ***D***, Experimental setup: light delivery in the south and west ChR2 arms, marked with dashed blue line. ***E***, Number of passes (left panel) and duration in seconds (right panel) counted in the ChR2 arms vs non-ChR2 arms during the pre-ChR2, ChR2- and post-ChR2 sessions. \*\**P* < 0.01. Error bars, mean ± s.e.m. ***F***, Number of passes through the ChR2- and non-ChR2 arms for the first 60 seconds of the post-ChR2 session. ***G***, Event related potentials (ERPs) recorded in dorsal CA1 from of 32 electrodes in a sample animal. Time 0 indicates the delivery of the onset of the BLA optogenetic stimulation. ***H***, Color-coded power spectrogram of hippocampal low-frequency oscillations (4 – 15Hz) during the photostimulation protocol. ***I***, Representative phase-locking value after BLA photostimulation for the observed data (blue) and for control shuffled data (green). ***J***, Comparison of the place cell’s spiking before and after the photostimulation onset for intra- and extra-field spikes. The pre/post normalised count represents decrease (positive) and increase (negative values) for the first 500- (left), 250- (middle) and 100ms (right) after the optogenetic protocol onset for the intra-field (light blue) and extra-field spikes (blue). \*\**P* < 0.01. Error bars, mean ± s.e.m. ***K***, ΔCOMa for place fields located outside the ChR2 arms (extra-field) and for place cells located inside the ChR2 arms (intra-field) from ChR2 recordings (blue) and control YFP recordings (green). \**P* < 0.05. Error bars, mean ± s.e.m.

**Figure 7.**
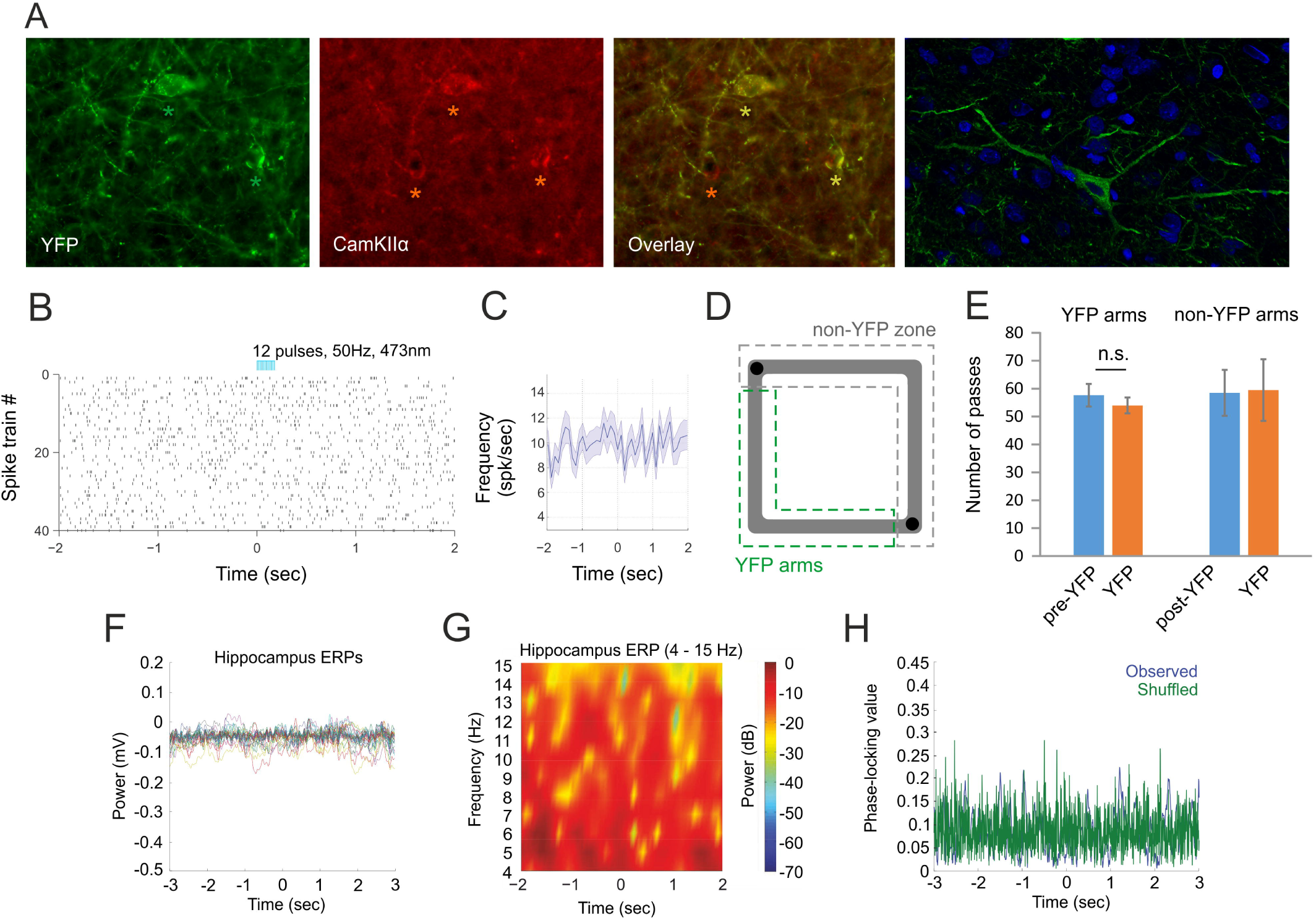
Photostimulation of control YFP-expressing BLA neurons. ***A***, YFP expression (left), anti-calcium/calmodulin-dependent protein kinase II alfa (anti-CamKIIα) staining (middle) and their overlay in BLA (right). The green asterisks show two YFP-expressing neurons, while the red asterisks show three CamKIIα-marked neurons. The confocal image on the far right shows YFP-expressing neurons (green) in BLA is histology with DAPI staining (blue). Raster plot from 40 repetitions ***B***, and ***C***, firing frequency of time-locked photostimulation of a BLA cell in control animals injected with AAV-YFP viral construct. Time 0 indicates the delivery of the first train of the stimulation protocol. ***D***, Experimental setup: light delivery in the south and west arms of the rectangular-shaped linear track (YFP arms, marked with dashed green line). No photostimulation was applied in the non-ChR2 zone (marked with dashed gray line). ***E***, Number of passes counted from control animals injected with AVV-YFP construct in the YFP arms vs non-YFP zone during the pre-YFP, YFP-, and post-YFP sessions. Error bars, mean ± s.e.m. ***F***, Event related potentials (ERPs) recorded in dorsal CA1 from of 32 electrodes in a sample control animal. Time 0 indicates the delivery of the onset of the BLA YFP photostimulation. ***G***, Color-coded power spectrogram of hippocampal low-frequency oscillations (4 – 15Hz) during the YFP photostimulation. ***H***, Representative phase-locking value after BLA YFP photostimulation for the observed data (blue) and for control shuffled data (green).

**Figure 8.**
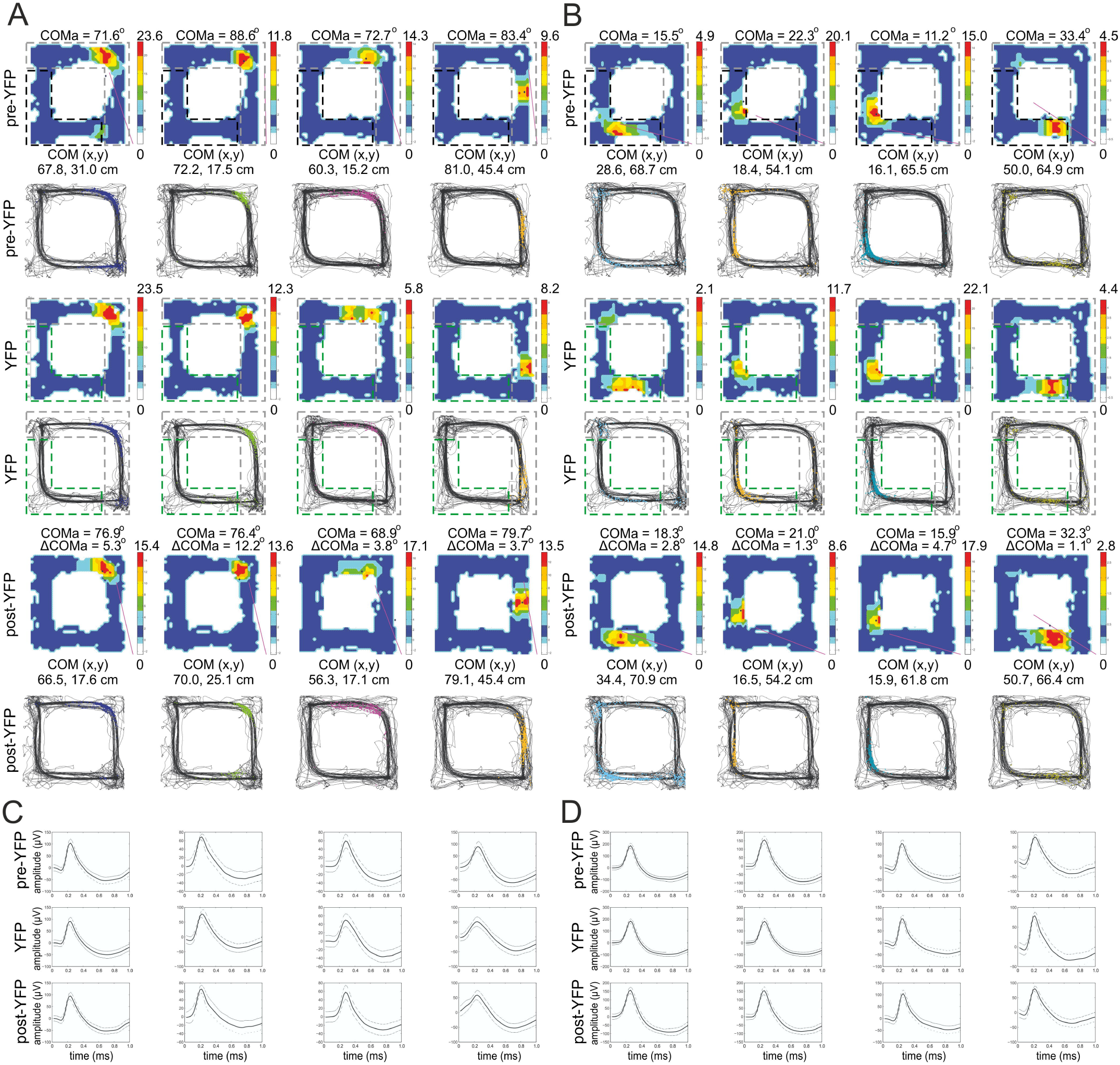
Control place fields during BLA photostimulation. ***A***, Four place fields from two sample animals with fields located outside the YFP arms during pre-YFP sessions (top), YFP (middle) and post- YFP sessions (bottom). For each session the upper panels show color-coded firing rate, while the lower panels show the animal trajectory with spikes, marked with colored dots. ***B***, Four place fields from the same two animals with fields located inside the YFP arms during pre-YFP sessions (top), YFP (middle) and post-YFP sessions (bottom). ***C***, Waveforms of the place cells shown in A, and ***D***, waveforms of the cells shown B, respectively. The solid line shows the average waveform shape; the dashed lines show the 1 SD confidence intervals.

### The extra-field spiking during BLA photostimulation predicts the center of mass shift

Place fields of the cells located in the non-ChR2 zone that spiked during the photostimulation session (Fig. 9) showed larger place field reconfiguration (Supplementary Movie 4), compared to the cells with little or no spiking activity (Fig. 10). We correlated the COM angle shift (ΔCOMa) to the ratio of the baseline over the TMT session place cells’ firing rate from the recorded spikes (ChR2 spiking ratio) for the place cells with fields located in the non-ChR2 zone (with extra-field spikes) as well as for the place cells located in the ChR2 arms (with intra-field spikes, Fig. 11A). The mean and the peak extra-field ChR2 spiking ratios significantly correlated to ΔCOMa (Fig. 11C, Pearson’s r = −0.429, *P* = 0.007; Fig. 11G, Pearson’s r = −0.426, *P* = 0.009, n = 38). Concurrently, the mean and the peak intra-field ChR2 spiking ratio showed weak non-significant correlation to ΔCOMa for the ChR2 arms (Fig. 11D, Pearson’s r = −0.261, *P* = 0.099; Fig. 11H, Pearson’s r = −0.187, *P* = 0.241, n = 41). No significant correlation to ΔCOMa was evident for the mean and peak firing rate of the intra- (Fig. 11E, Pearson’s r = 0.092, *P* = 0.530, n = 49; Fig. 11I, Pearson’s r = 0.066, *P* = 0.651, n = 49) and extra-field spikes (Fig. 11F, Pearson’s r = 0.099, *P* = 0.529, n = 42; Fig. 11J, Pearson’s r = 0.032, *P* = 0.842, n = 42) for the non-ChR2 zone. Similarly to the TMT protocol, the BLA photostimulation evoked significant correlation between the spiking ratio and ΔCOMa only for the ChR2 arms and only for the extra-field spiking cells. These data validate the hypothesis that aversion-evoked place field reconfiguration is mediated by BLA activation.

**Figure 9.**
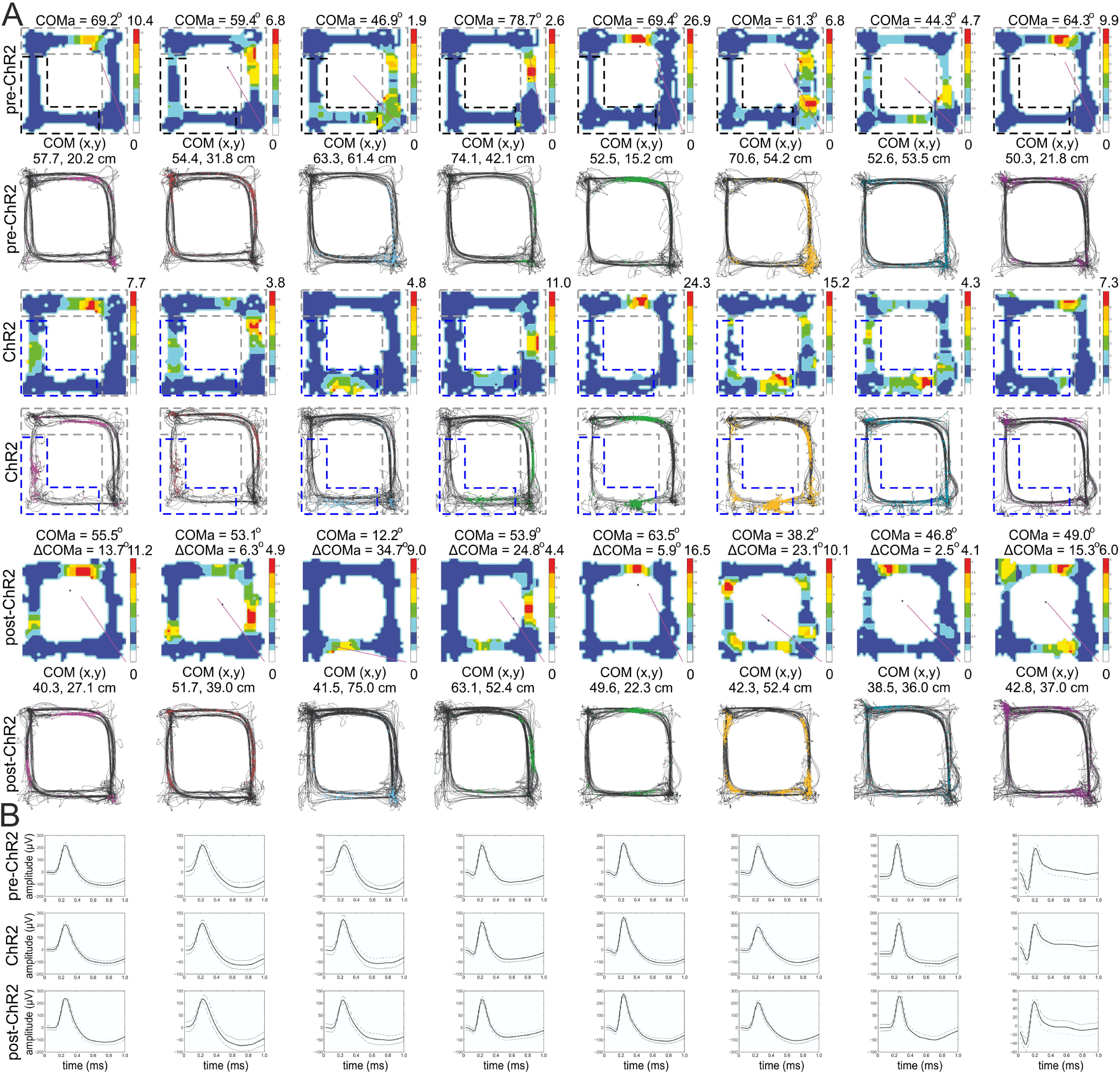
Place fields of cells with high extra-field spiking during BLA photostimulation. ***A***, Eight place cells from four rats, with fields located outside the ChR2 arms, with expressed extra-field spiking activity during the light delivery. Top panels: pre-ChR2 session, middle panels: ChR2 session, and bottom panels: post-ChR2 session. For each session the upper panels show color-coded firing rate, while the lower panels show the animal trajectory with spikes, marked with colored dots. ***B***, Waveforms of the place cells above, recorded from the pre-ChR2 (top row), ChR2 (middle row) and post-ChR2 session (bottom row). The solid line shows the average waveform shape; the dashed lines show the 1 SD confidence intervals.

**Figure 10.**
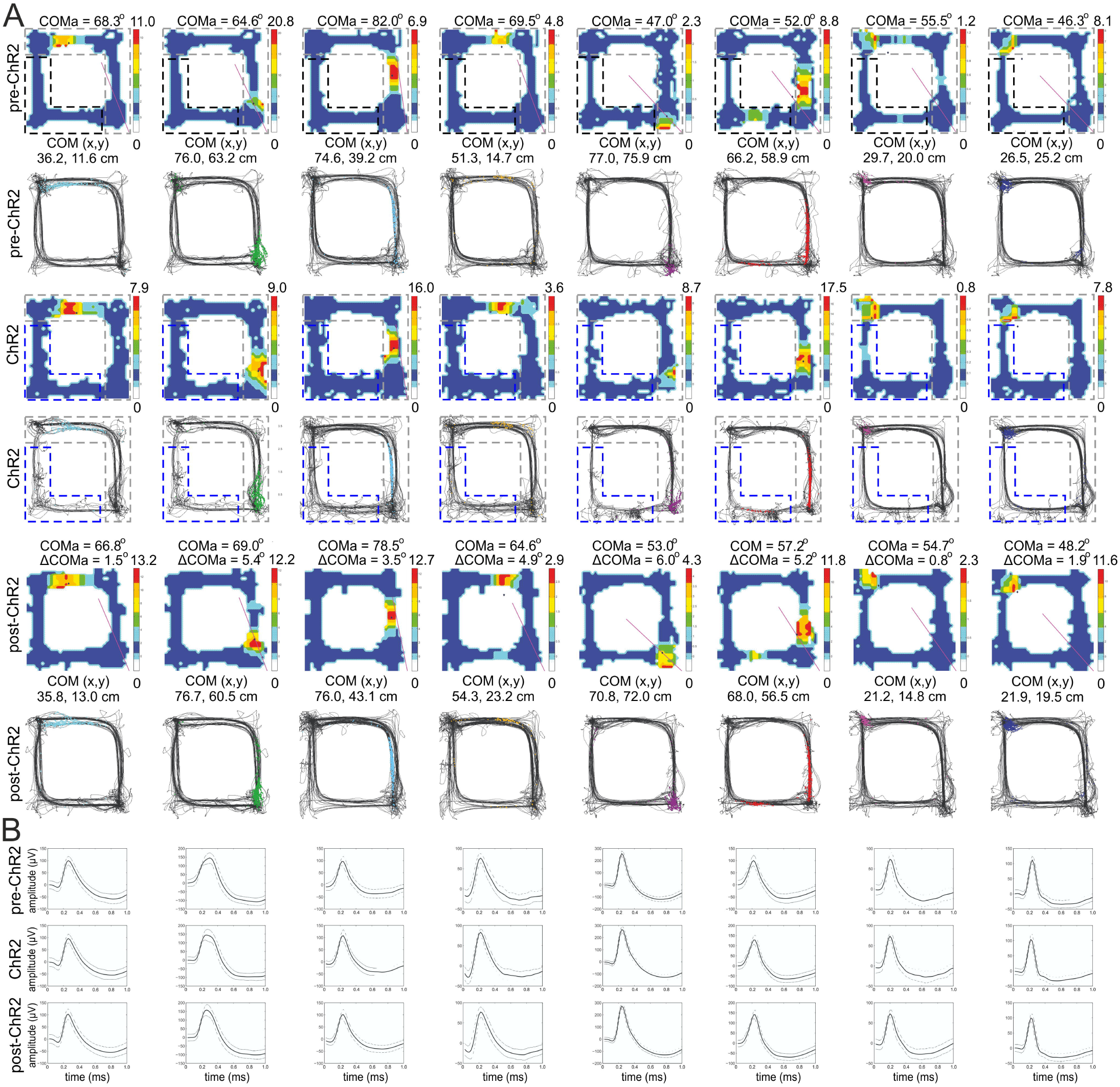
Place fields of cells with low extra-field spiking during BLA photostimulation. ***A***, Eight place cells from the same four rats, with fields located outside the ChR2 arms, with little or no extra-field spiking activity during the light delivery. For each session the upper panels show color-coded firing rate, while the lower panels show the animal trajectory with spikes, marked with colored dots. ***B***, Waveforms of the place cells above, recorded from the pre-ChR2 (top row), ChR2 (middle row) and post-ChR2 session (bottom row). The solid line shows the average waveform shape; the dashed lines show the 1 SD confidence intervals.

**Figure 11.**
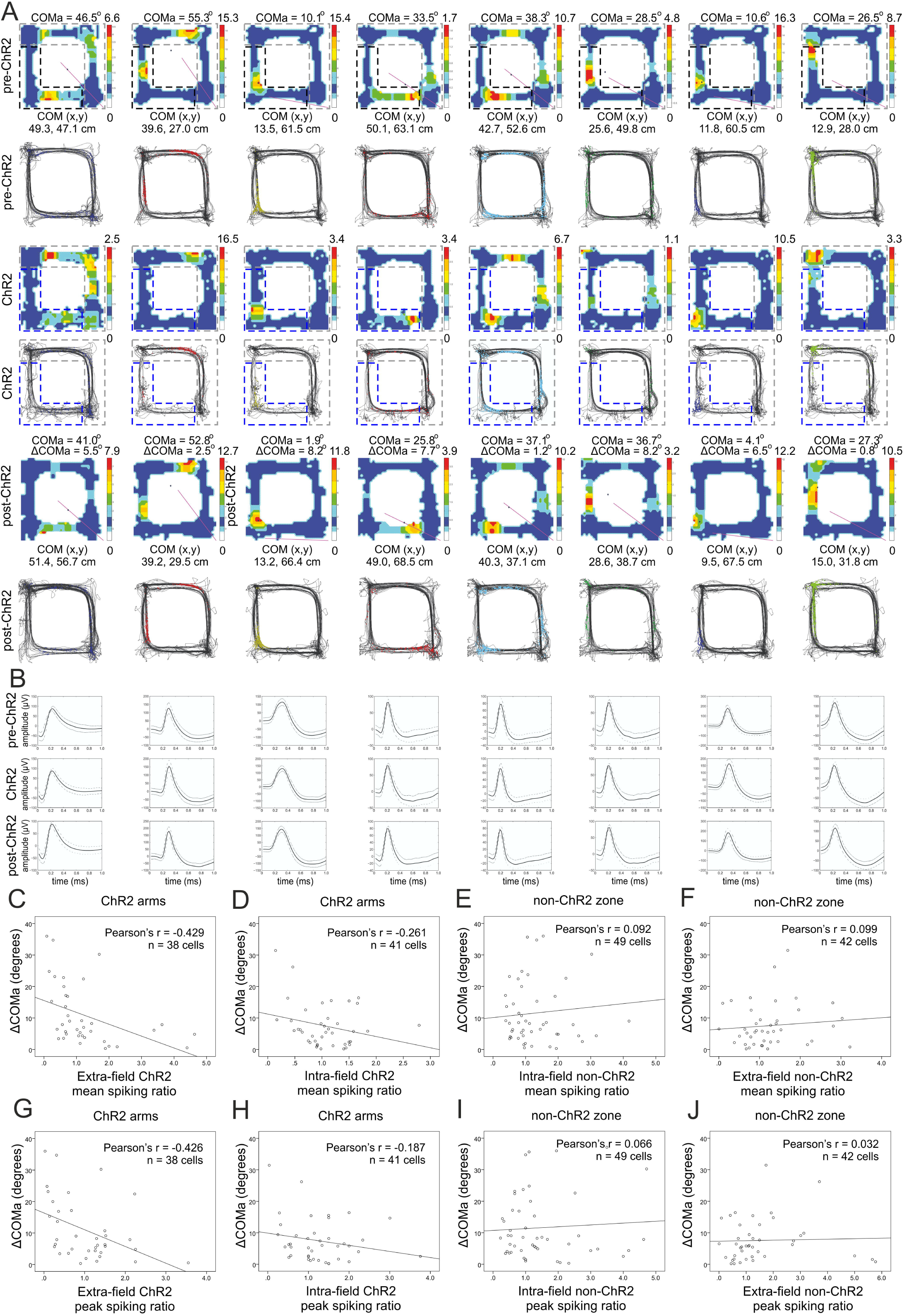
BLA-triggered hippocampal spiking increase correlates to the place field center of mass shift. ***A***, Eight place cells from four rats, with fields located inside the ChR2 arms, with intra-field spiking during the light delivery. Top panels: pre-ChR2 session, middle panels: ChR2 session, and bottom panels: post-ChR2 session. For each session the upper panels show color-coded firing rate, while the lower panels show the animal trajectory with spikes, marked with colored dots. ***B***, Waveforms of the place cells above, recorded from the pre-ChR2 (top row), ChR2 (middle row) and post-ChR2 session (bottom row). The solid line shows the average waveform shape; the dashed lines show the 1 SD confidence intervals. ***C***, Correlation between ΔCOMa and ChR2 mean spiking ratio for the extra-field spikes of the place cells located outside the ChR2 arms, Supplementary Table 11. ***D***, Correlation between ΔCOMa and ChR2 mean spiking ratio for the intra-field spikes of the place cells located inside the ChR2 arms, Supplementary Table 12. ***E***, Correlation between ΔCOMa and non-ChR2 mean spiking ratio for the intra-field spikes of the place cells located inside the non-ChR2 zone, Supplementary Table 13. ***F***, Correlation between ΔCOMa and non-ChR2 mean spiking ratio for the extra-field spikes of the place cells located outside the non-ChR2 zone, Supplementary Table 14. ***G***, Correlation between ΔCOMa and ChR2 peak spiking ratio for the extra-field spikes of the place cells located outside the ChR2 arms, Supplementary Table 11. ***H***, Correlation between ΔCOMa and ChR2 peak spiking ratio for the intra-field spikes of the place cells located inside the ChR2 arms, Supplementary Table 12. ***I***, Correlation between ΔCOMa and non-ChR2 peak spiking ratio for the intra-field spikes of the place cells located inside the non-ChR2 zone, Supplementary Table 13. ***J***, Correlation between ΔCOMa and non-ChR2 peak spiking ratio for the extra-field spikes of the place cells located outside the non-ChR2 zone, Supplementary Table 14.

### The unidirectional field shift after BLA photostimulation is similar to the TMT-induced remapping

We compared the spiking parameters of the place fields between pre-YFP session (Fig. 12A,B left panels) and post-YFP session (Fig. 12A,B right panels), as well as pre-ChR2 session (Fig. 5C,D left panels) and post-ChR2 session (Fig. 12C,D right panels) of the BLA-photostimulated group. There was no significant difference in the change of the mean firing rate between the YFP and ChR2 groups for the clockwise (Fig. 12E, one-way ANOVA with Bonferroni post-hoc correction, n = 6, F_(1,122)_ = 0.269, *P* = 0.847) and counter-clockwise groups (Fig. 12H, one-way ANOVA with Bonferroni post-hoc correction, n = 6, F_(1,112)_ = 0.278, *P* = 0.810). No significance was evident for the change of the peak firing rate between both groups for the clockwise (Fig. 12F, one-way ANOVA with Bonferroni post-hoc correction, n = 6, F_(1,122)_ = 0.266, *P* = 0.850) and for counter-clockwise fields (Fig. 12I, one-way ANOVA with Bonferroni post-hoc correction, n = 6, F_(1,112)_ = 0.126, *P* = 0.816). The linearized ΔCOM was 5.66 ± 1.1 cm for the clockwise- (Fig. 5G) and 5.61 ± 1.1 cm for the counter-clockwise fields (Fig. 5J) in the YFP group, while light delivery to BLA induced ΔCOM of 24.60 ± 4.6 cm for clockwise- and 24.10 ± 3.0 cm for (counter-clockwise fields one-way ANOVA with Bonferroni post-hoc correction, n = 6, F_(1,122)_ = 12.01, *P* < 0.001; counter-clockwise fields one-way ANOVA with Bonferroni post-hoc correction, n = 6, F_(1,112)_ = 11.07, *P* < 0.001). These data show that similarly to TMT, BLA excitation results in strong field but insignificant rate remapping for both directions of navigation.

**Figure 12.**
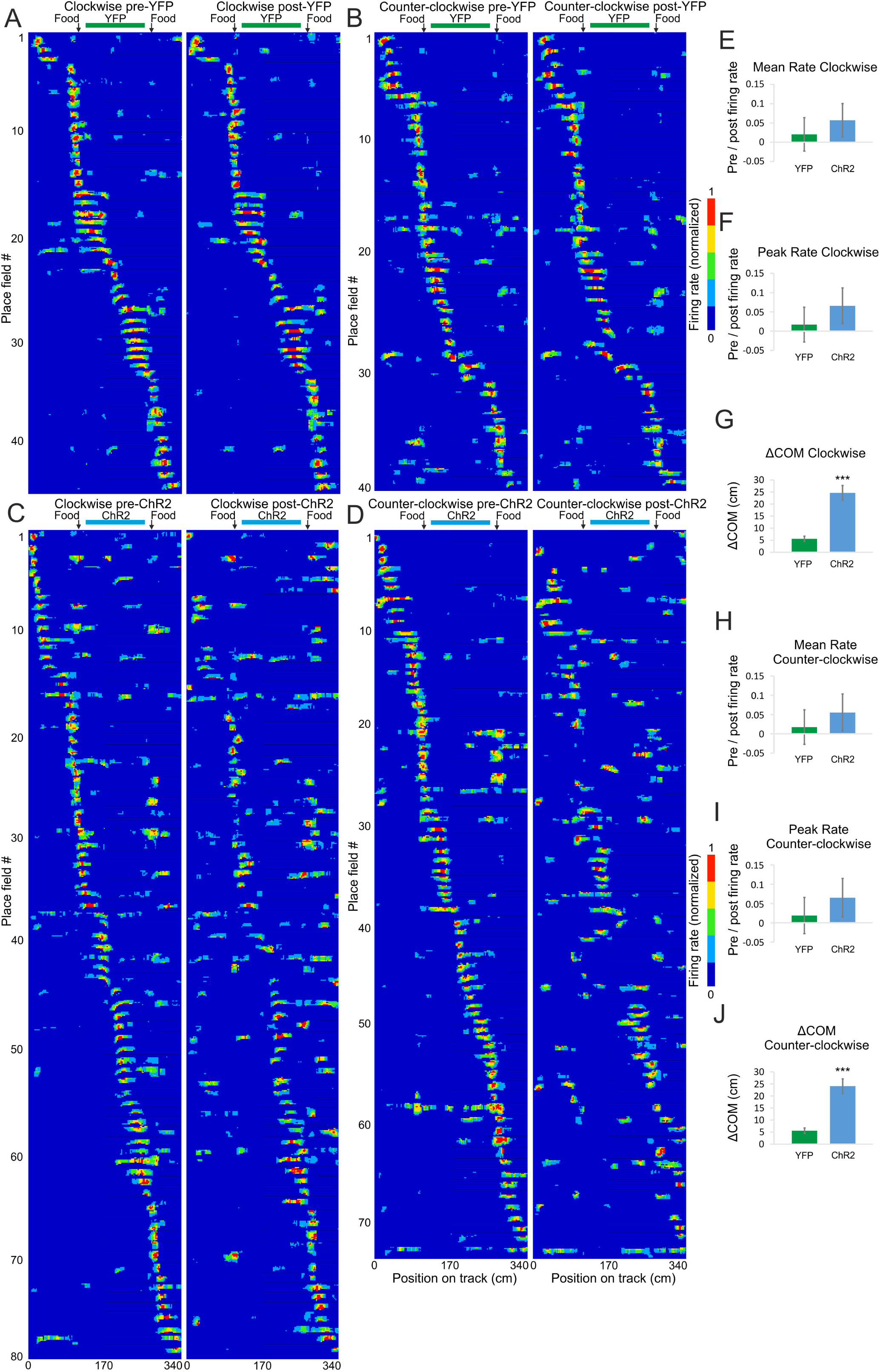
Unidirectional field remapping after BLA photostimulation. ***A***, Color-coded linearized map showing location of CA1 place fields before (left panel) and after (right panel) light delivery session to the YFP group of rats for clockwise, Supplementary Table 15 and ***B***, counter-clockwise direction of movement, Supplementary Table 16. Each line shows activity of one place cell (84 datasets in total from 69 place cells). The horizontal green bar indicates the light delivery YFP zone during the exposure session, while black vertical arrows indicate the location of the food delivery. ***C***, Color-coded linearized map showing location of CA1 place fields before (left panel) and after (right panel) light delivery session to the ChR2 group of rats for clockwise, Supplementary Table 17 and ***D***, counter-clockwise direction of movement, Supplementary Table 18. Each line shows activity of one place cell (154 datasets in total from 79 place cells). The horizontal blue bar indicates the light delivery ChR2 zone during the exposure session, while black vertical arrows indicate the location of the food delivery. ***E***, Comparison of the place field mean spiking before and after BLA photostimulation. The pre/post normalised count represents decrease (positive) and increase (negative values) for the mean firing rate of clockwise fields for YFP and ChR2 groups. Error bars, mean ± s.e.m. ***F***, Comparison of the place field peak spiking before and after BLA photostimulation of clockwise fields for the same groups. Error bars, mean ± s.e.m. *G*, Center of mass shift (ΔCOM) after BLA photostimulation of clockwise fields for YFP and ChR2 groups. Error bars, mean ± s.e.m. \*\*\**P* < 0.001. ***H***, Comparison of the place field mean spiking before and after BLA photostimulation for counter-clockwise fields. Error bars, mean ± s.e.m. ***I***, Comparison of the place field peak spiking before and after BLA photostimulation for counter-clockwise fields. Error bars, mean ± s.e.m. ***J***, Center of mass shift (ΔCOM) after BLA photostimulation for counter-clockwise fields. Error bars, mean ± s.e.m. \*\*\**P* < 0.001.

## Discussion

In this study, we demonstrated the reconfiguration pattern of hippocampal place cells evoked by aversive episodes. We report that the place fields located outside the area of aversion express higher propensity for center of mass shift, compared to the fields located with the arms of the track associated with TMT-induced aversion. Our findings show for the first time that the degree of center of mass shift correlates with the firing rate of the extra-field spiking during the aversion. Optogenetic stimulation of extra-field spikes resulted in the same reconfiguration pattern with the exposure to innately aversive odor.

### Beta amplitude increase displays the spatial location of aversive odor perception

The hippocampus is proposed to provide a contextual framework for the encoding of emotional events (Leutgeb et al., 2005). Recent data demonstrate that hippocampal spatial representation is not static but alters in response to non-spatial rewarding stimuli (Mamad et al., 2017). Hippocampal place cells remap also after exposure to stressful or fearful events (Moita et al., 2004; Wang et al., 2012; Kim et al., 2015; Wang et al., 2015). This remapping is a mechanism for the formation of new hippocampal engrams that store the association of spatial context and fearful experience (Ramirez et al., 2013; Tonegawa et al., 2015). Although a population of place cells remap after exposure to stressful or fearful events (Moita et al., 2004; Kim et al., 2015), there is no clear explanation why some neurons alter their place fields to encode fearful experience, while others preserve stable fields. The application of innate aversive trimethylthiazoline (TMT) allowed us to study the transformation of hippocampal population code from indifferent to negative context value for a single environment. The advantage of TMT is the absence of learning curve required for associative fear conditioning (Kobayakawa et al., 2007) and the absence of visible or tactile features of the stressor that might affect the stability of the recorded place cells (Jeffery and O’Keefe, 1999). Beta rhythm (15 – 40 Hz) is a characteristic oscillation in the olfactory bulb during odorant perception (Lowry and Kay, 2007; Lepousez and Lledo, 2013). During olfactory-driven behavior beta oscillations sustained long-range interactions between distant brain structures, including the hippocampus (Martin et al., 2007). We measured beta oscillations to identify in which arms of the track the animals perceive the aversive odorant. This approach allowed us to distinguish whether TMT odor was processed by the hippocampal formation of the animals only for particular arms or for the entire rectangular-shaped linear track. Entorhinal–hippocampal coupling was observed specifically in the 20 – 40 Hz frequency band as rats learned to use an odor cue to guide navigational behavior (Igarashi et al., 2014). The essence of beta rhythm in the processing of aversive stimuli across the limbic circuitry was demonstrated with the finding that powerful beta activity predicted the behavioral expression of conditioned odor aversion (Chapuis et al., 2009). Similarly, we showed that beta amplitude increased during the TMT sessions, particularly in the TMT arms but not in the rest of the track. We revealed here that beta rhythm varied as a function of speed: the frequency of beta decreased, while the amplitude increased with the augmentation of the whole body motion of the animal. The frequency dependence on the speed of locomotion is not characteristic only for the beta frequency band but also for slower and faster local field oscillations. Increase of gamma frequency was linked to faster running speed in rats (Ahmed and Mehta, 2012) and this phenomenon was proposed to preserve the spatial specificity of place cells at different running speeds (Ahmed and Mehta, 2012). Further research is needed to clarify the source of the currents underpinning beta. Particularly intriguing is the open question of how the hippocampal pyramidal cells discharge in relation to beta phase during the onset of aversive experience. We need to elucidate the temporal link between beta and theta, as well beta and gamma during the perception of aversive stimuli.

### Aversion experience reconfigures hippocampal place fields

After we identified the TMT arms as a section of the maze where the animals perceived the aversive stimulus, we compared the hippocampal spatial representation across the track axis, dissociating the TMT arms against the opposing arms. The reconfiguration of individual place fields is best evaluated by the center of mass difference (Knierim, 2002). We found that place fields from animals exposed to aversive TMT evoked higher center of mass change compared to control animals. Exposure to predator odor produces only partial remapping and one possibility is that the cells remaining stable encode visuospatial information, while the remapping cells are sensitive to other contextual cues such as olfactory information and/or emotional valence (Wang et al., 2012). Another possibility is that spatial or temporal proximity of the neuronal activity to the perception of the aversive episode determines which place fields reconfigure. To show that the spatial proximity to the olfactory stimulus was essential component of the field plasticity we compared the correlation between the change of the center of mass angle (ΔCOMa) and TMT spiking ratio within the TMT arms versus outside the TMT arms (non-TMT zone). We used ΔCOMa as analytical tool to evaluate the displacement of place fields in relation to aversive event and to measure the change of their spatial proximity to the TMT arms. Surprisingly, we found that the first group (including intra-field spikes) showed no significant correlation, while the second group (including only extra-field spikes) demonstrated high correlation of the spiking activity with the degree of center of mass shift. No significant correlation was evident between ΔCOMa and intra- or extra-field spiking ratio for the non-TMT zone of the track. Although information on the contribution of extra-field spikes to spatial navigation is lacking in the literature, we know that their activity is crucially involved in hippocampus-dependent learning (Ferguson et al., 2011). The significance of the extra-field spikes is largely associated to the sharp-wave ripple replay of recently navigated locations and experiences (Wu et al., 2017), which are important in learning and memory consolidation during inactive behavioural state (Johnson et al., 2009). However, extra-field spikes from place cells have been shown to occur during active navigation (Johnson and Redish, 2007; Epsztein et al., 2011). Here, we present evidence that the extra-field activity encodes salient stimuli and if it is potent enough to evoke place field plasticity related to experience-dependent learning. The TMT protocol allows for instant association between the aversive episode and spatial location, which is easily detected by the avoidance behaviour and validated by the beta rhythm amplitude. The advantage of a snapshot aversion episode is the robust change of the place cells spiking inside and outside their place fields.

Not only aversion but also reward and novelty salient stimuli can trigger biased field distribution (Fyhn et al., 2007). Our experimental design included two feeding locations. Therefore, these may have contributed to the observed remapping. The delivery of sugar pellets ensured continuous navigation for 12 minutes for each recording session. To reduce the effect of the reward on the place field remapping we habituated the rats to the feeding locations. The animals were consistently rewarded throughout the baseline (pre-) recordings, TMT/ethanol or YFP/ChR2 recordings and subsequent (post-) recordings for all groups of animals. The unidirectional field analysis revealed that TMT or ChR2 groups of rats revealed significantly greater shift in COM (increased ΔCOM) compared to the ethanol or YFP groups, respectively. The observed field plasticity was field but not rate remapping. However, this result does not rule out the possibility that this plasticity results from a change in reward value due to TMT exposure rather than the mere effect of conditioned aversion. A study in behaving rats showed that a decrease in lever pressing for food in the presence of a conditioned tone also led to changes in firing rate in areas associated with fear learning (Sotres-Bayon et al., 2012). Therefore, the perception of known reward value may change due to conflict between aversion and reward. We must acknowledge the likelihood that the COM shift observed in the non-TMT (non-ChR2) arms may be the result of conflict between the rewarding and aversive stimuli. Another factor that can affect place field variability is the speed of navigation. Although the post-TMT(ChR2) sessions were characterized with reduced navigation during the first two minutes, the animals navigation recovered for the remaining 10 minutes. This is of sufficient duration for the reliable formation of stable fields (Frank et al., 2004). Concurrently, we excluded spikes that occurred during epochs with running speeds below 5 cm/s (Alme et al., 2014).

### Amygdala mediates aversion-induced shift of the place field center of mass

The amygdala is a key structure in the acquisition of aversive experience (LeDoux, 2000) and optogenetic stimulation of BLA mediates associative fear learning (Johansen et al., 2010; Klavir et al., 2017). The amygdala sets the emotional valence of sensory stimuli (Phelps and LeDoux, 2005; Moriceau et al., 2006) and the BLA circuitry is particularly involved in the odor-evoked fear conditioning (Wallace and Rosen, 2001; Anderson et al., 2003; de Araujo et al., 2003). The amygdala is critical for stress-induced modulation of hippocampal synaptic plasticity and hippocampus-dependent learning (Kim et al., 2001; Vouimba and Richter-Levin, 2005). Aversion-triggered place cell remapping is blocked by amygdala inactivation (Donzis et al., 2013; Kim et al., 2015), while BLA stimulation decreases the stability of CA1 place fields (Kim et al., 2012). Despite recent advances in manipulating engrams (Cai et al., 2016; Rashid et al., 2016), there is no agreement if the place cells remap randomly or their reconfiguration depends on the location of the aversive stimulus perception. We found that the BLA-triggered field plasticity pattern was equivalent to the TMT-induced field reconfiguration, namely, the correlation between the ChR2 spiking ratio and ΔCOMa was significant only for place cells located outside the ChR2 arms. The non-ChR2 zone included non-ChR2 arms and the feeding zones. Elimination of the feeding zones from the data analyses would have excluded spikes from intra- or extra-field activity, affecting the reliability of the place field center of mass calculation and the accuracy of field plasticity identification. The observed field reconfiguration pattern reveals that the increase of extra-field but not intra-field spiking during aversive episode predicts the change in the preferred firing location. Such response can emerge after a small, spatially uniform depolarization (Rickgauer et al., 2014) of the spatially untuned somatic membrane potential of inactive place cell leads to the sudden and reversible emergence of a spatially tuned subthreshold response and novel place field formation (Lee et al., 2012; Bittner et al., 2015). This phenomenon is proposed as a key mechanism for the formation of hippocampal memory representations (Lee et al., 2012). Our data complement these findings by showing that BLA stimulation increases only the extra-field but not the intra-field spiking of the CA1 place cells. It is very likely that extra-field spikes might by highly-susceptive to hippocampal inputs and these spikes mediate spike-timed synaptic plasticity that results in relocation of the place fields’ center of mass. Therefore, our data analyses were not restricted only to the main place field, but included also the identification of the extra-field spikes. Early studies showed that amygdala can regulate the induction of hippocampal synaptic plasticity, where the population spike long-term potentiation in the dentate gyrus was attenuated by lesion of the BLA (Ikegaya et al., 1994) or by pharmacological inactivation of BLA (Ikegaya et al., 1995). Priming the BLA inputs 30 seconds prior to performant path stimulation resulted in the facilitation of the excitatory post-synaptic potentials in the dorsal hippocampus (Akirav and Richter-Levin, 1999). Thus, BLA modulates the synaptic plasticity within the hippocampal formation where the amygdala and the hippocampus acts synergistically to form long-term memories of emotional events. The dual activation of the amygdala and the hippocampus and the cross-talk between is proposed to provide contextual information to emotionally based memories (Richter-Levin and Akirav, 2000). The dorsal hippocampus is believed to associate context and fear memories (Ryan et al., 2015; Tonegawa et al., 2015) and our data suggest that this process can be detected in the spatial firing patterns of the place cells, which shift their center of mass towards the aversive section of the environment. While our experimental design involves brief aversion retrieval (during the first minute of the post-TMT/ChR2 sessions) followed by aversion extinction, the observed field reconfiguration may be specific for the extinction phase. The place field remapping pattern between the retrieval and extinction may differ (Wang et al., 2015), however, the long-term measurement of place cell activity during aversion retrieval is challenging due to the risk of navigation undersampling which is a reason for incomplete formation of place fields *per se* and invalidates the evaluation of their properties (Hok et al., 2012; Navratilova et al., 2012).

### Contextual signalling across the hippocampal formation

Although the physiology of dorsal hippocampus is regulated by the aversive amygdalar signals it is still unclear which is the most direct anatomical route from BLA to dorsal hippocampus. The most substantial projection to the hippocampus originates in the basal nucleus and the caudomedial portion of BLA projects heavily to the stratum oriens and stratum radiatum of hippocampal CA3 and CA1 with predominant innervation of the ventral hippocampus (Pikkarainen et al., 1999). Thus, the BLA-dorsal hippocampus signal transmission most likely follows indirect polysynaptic route via the ventral hippocampus (Amaral and Witter, 1989). Lesions of the longitudinal hippocampal pathways demonstrated the functional significance of ventro-dorsal projections in spatial memory formation (Steffenach et al., 2002). Ventral hippocampus is known also to mediate contextual conditioning where ventral hippocampal lesions disrupt contextual freezing (Maren and Holt, 2004; Orsini et al., 2011; Kim and Cho, 2017), but see (Huff et al., 2016). Furthermore, studies in vivo have shown that cells in the ventral region provide contextual information (Komorowski et al., 2013) and ventral cells respond to odors much more strongly than dorsal cells (Keinath et al., 2014). This line of research reveals the role of ventral hippocampus in fear and contextual conditioning and suggests that aversive signals may propagate across the hippocampal longitudinal axis towards dorsal hippocampus that mediates spatial learning. We found that BLA photostimulation triggered potent oscillatory response in dorsal hippocampus represented by ERPs synchronization after the light pulse delivery and increased power in the range of 5 – 8 Hz. The firing of the hippocampal neurons also changed with higher occurrence of extra-field spikes 100 ms after the photostimulation, although we did not find hippocampal spikes that were directly entrained by the light pulses. The electrophysiological data presented in our manuscript report functional relation between the amygdala and dorsal hippocampus, but our histological data were insufficient to conclude whether this effect was mediated by the major indirect ventral projections or by sparse direct dorsal projections. Therefore, we consider that the indirect ventral pathway as the most likely anatomical route that mediated the observed amygadalo-hippocampal signalling. Spike-timed depolarization may underline not only the encoding of aversive but also of rewarding or salient stimuli mediated by the subcortical or entorhinal projections to hippocampus (Harvey et al., 2009).

We report here that BLA-induced field remapping follows the place field plasticity patterns after aversive experience, evoked with exposure to TMT. Our data should be also considered in the context of two possible conditions: 1) the remapping patterns may occur not only as a result of the TMT or ChR2 protocol but also due to conflict between the rewarding and aversive stimuli, and 2) the plasticity of dorsal hippocampal place cells may be mediated via different indirect pathways arising from BLA. We propose that this pattern of field reconfiguration serves as a universal mechanism for the generation of multiple context-dependent representations by different salient stimuli, where the animal’s behavior is guided by the contextual valence of previous experience.

## Acknowledgements

This work was supported by Science Foundation Ireland, the Health Research Board and the Wellcome Trust under Biomedical Research Partnership with grant #099926/Z/12/Z to Marian Tsanov.

